# Single-cell-guided identification of logic-gated antigen combinations for designing effective and safe CAR therapy

**DOI:** 10.1101/2025.03.19.644074

**Authors:** Sanna Madan, Tian-Gen Chang, Alexandra R. Harris, Huaitian Liu, Andrew Martinez, Saugato Rahman Dhruba, Binbin Wang, Padma Sheila Rajagopal, Sanju Sinha, Aravind Srinivasan, Simon R. V. Knott, Shahin Sayed, Francis Makokha, Chi-Ping Day, Gretchen L. Gierach, Stefan Ambs, Alejandro A. Schäffer, Eytan Ruppin

## Abstract

Chimeric antigen receptor (CAR) T-cell therapy has revolutionized the treatment of hematological malignancies. However, its application in solid tumors remains limited because single targets are unlikely to suffice due to tumor antigen heterogeneity and off-tumor toxicities. To overcome these obstacles, we developed *LogiCAR designer*, a computational approach that utilizes single-cell transcriptomics data from patient tumors to systematically identify the cancer-specific antigen circuits with logic gates (“AND,” “OR,” and “NOT”) that target the majority of cancer cells in a tumor while sparing normal cells and tissues as much as possible. *LogiCAR designer* efficiently scales to higher-order antigen combinations involving up to five genes. Applied to a large-scale dataset encompassing approximately 2 million cells (including > 620k tumor cells) from 342 clinical patient samples across all major breast cancer subtypes, *LogiCAR designer* identified antigen circuits with enhanced tumor-targeting efficacy and improved safety profiles compared to both previously reported circuits and single-target therapies in clinical trials. However, even these optimized shared circuits still proved insufficient for some patients. We hence systematically studied *LogiCAR designer*’s ability to identify highly effective CAR circuits that are individualized to each patient. Remarkably, such personalized CAR circuits provide estimated tumor-targeting efficacy tantamount to complete response in 76% of patients and partial response for all patients. Taken together, this analysis is the first systematic quantification of the efficacy and safety of all possible CAR circuits, showing that: **(a)** the quality of existing solutions leaves much to be desired; **(b)** the ability of shared circuits optimized across many patients is moderate, and finally, **(c)** individually tailored circuits offer significantly higher tumor-targeting efficacies for patients. *LogiCAR designer* offers a rigorous, data-driven way to facilitate the rational design of safe and effective CAR-based immunotherapies for cancer.

**Statement of Significance:**

⍰ Development of a computational approach that efficiently identifies logic-gated CAR target combinations, called *circuits,* from single-cell transcriptomics, addressing a critical unmet clinical need.
⍰ Application to the largest ensemble of breast cancer datasets to date, comprising ∼2 million cells (> 620k tumor cells) from 17 clinical cohorts, to identify CAR circuits predicted to be effective.
⍰ Comprehensive safety profiling of candidate circuits spanning major tissues at both RNA and protein levels.
⍰ Logic-gated CAR circuits generated by our pipeline address tumor heterogeneity and achieve efficacy and safety scores that surpass clinical trial and previously computationally identified circuits.
⍰ Individualized rational CAR design offers a transformative approach to deliver precision-engineered CAR therapies with unprecedented efficacy.

## Introduction

Chimeric antigen receptor (CAR) T-cell therapies have revolutionized cancer treatment, particularly for cancers of B cell lineages. All six CAR T-cell therapies approved by the U.S. Food and Drug Administration (FDA) have indications for B cell-lineage cancers, including B cell lymphomas and multiple myeloma ^1^. These therapies eliminate all cells expressing the B-cell lineage markers (e.g., CD19 or TNFRSF17) on the cell surface, and the side effects can be managed with intravenous immunoglobulin ^1,2^. In CAR-T therapy, T cells are extracted from the patient’s blood, engineered to express the synthetic CAR that recognizes tumor-associated antigen(s), and reinfused into the patient ^1^. While highly successful for B cell cancers, CAR-T faces challenges in other tumor types where complete elimination of the cell lineage would be fatal ^3^.

A major challenge is the heterogeneity of antigens expressed on cancer cell surfaces – targeting only one antigen may not result in the CAR T cells recognizing sufficiently many cancer cells in a patient to be of clinical benefit ^4,5^. In some types of cancers, such as triple negative breast cancer (TNBC), there are no obvious single tumor-associated antigens to target. Additionally, antigens loosely considered to be cancer-specific often appear on healthy cell surfaces as well. Addressing these challenges has led us to define a formal notion of optimal antigen selection and/or counter-selection and to design a strategy to search for antigen circuits that are best possible according to that definition. Our definition of optimality can be applied in an individualized manner to single patients or to entire cohorts by averaging over all patients.

In clinical trials to date, targeting single antigens in solid tumors and non-B-cell hematological malignancies has shown low efficacy and off-tumor side effects. To mitigate this, several groups have developed systems for logic-gated targeting of multiple antigens ^6–9^, in which CARs would target multiple genes to control T cell activity through Boolean logic gates: AND (&), OR (|), and NOT (!). For example, ‘G1 | G2’-activating CAR T cells target cells expressing either G1 or G2; ‘G1 | (G2 & G3)’ is for targeting cells expressing G1 or both G2 and G3; and ‘G1 | (G2 & !G3)’ is for targeting cells expressing G1 or G2 but not G3. Such control is made possible by combining multiple extracellular antigen binding domains and/or intracellular signaling domains in CAR. Logic-gated CAR NK cells ^10^ and T cells ^6^ have already been shown to address intra-tumor heterogeneity and reduce off-tumor effects.

Previous bioinformatic studies have considered two related problems: (1) identifying the best single targets and (2) identifying the best combinations. Either problem could be studied using bulk transcriptomics or single-cell transcriptomics. For example, bulk transcriptomics was used to find both single targets ^11^ and target combinations ^12,13^ previously. Single-cell transcriptomics of patient tumors were used to identify single targets for either head and neck cancer or acute myeloid leukemia ^14,15^. Kwon and colleagues used single-cell transcriptomics to find circuits of up to two targets ^16^. We previously developed the software MadHitter that uses single-cell transcriptomics to answer what is the minimum number of targets with logical ‘OR’ gating needed to meet a tradeoff between targeting a certain threshold proportion of tumor cells while sparing a different threshold proportion of non-tumor normal cells ^17^. Bulk expression data can obscure true cancer-cell-specific signals due to its mixed cell type content, and Dannenfelser *et al.*’s bulk transcriptomics analysis ^12^ was limited to AND and NOT gating. The study of Dannenfelser and colleagues ^12^ included some wet lab validation and the top single target prediction of Ahmadi and colleagues, *PTPRZ1* in brain cancer, was subsequently validated *in vivo* by Martinez Bedoya and colleagues ^18^, demonstrating that bioinformatic derivation of targets has translational potential. Fully exploring the range of Boolean gating with AND, OR, and NOT logic, and circuits of three or more surface targets, has remained a fundamental open challenge, to be carried out and benchmarked against previous approaches.

Screening all possible circuits of three or more cell surface protein-encoding genes across all cells is computationally intensive if the number of cells is large. To address this gap, we developed a genetic algorithm-based approach, termed *LogiCAR designer*, that analyzes single-cell transcriptomics from patient tumors and healthy human tissues as its input. *LogiCAR designer* then identifies optimal logic gates of surface antigens that are specifically activated in cancer cells and are only minimally active in normal, healthy cells across healthy tissues to ensure treatment safety. We demonstrate that *LogiCAR designer* successfully identifies gene triplet circuits that enhance selectivity and safety, outperforming clinical trial-tested CAR antigen targets. *LogiCAR designer* scales efficiently to circuits of larger size and performs robustly across diverse parameter settings. *LogiCAR designer* is applied to analyze an extensive collection of breast cancer (BRCA) single-cell and single-nucleus datasets on an unprecedented scale, including a new, unpublished resource spanning numerous clinical subtypes and ethnicities. The resulting newly identified circuits consistently exhibited superior performance versus existing solutions. Notably, this includes logic-gated circuits targeting TNBC, the most resistant BRCA subtype, which lacks established targets. More generally, *LogiCAR designer* is a promising approach toward designing more selective and safer next-generation CAR therapies for any cancer type of interest.

## Results

### Overview of this study

This study aimed to identify effective and safe CAR circuits for cancer treatment, including single-gene targets and, importantly, logic-gated combinations (**Fig. 1A**). Henceforth, we use the electrical engineering term *circuit* to describe a formula in which the variables are *specific* genes, and the genes are combined via AND, OR, NOT gates; in some contexts, ‘circuit’ will also refer to single-gene CARs that have no gates. We use the term *design* or *logic-gate design* to refer to the combinatorial structure of the gates without the gene variables assigned; for example, ‘((gene1 | gene2) & gene3)’ is three-gene, two-gate design, where ‘geneN’ does not refer to any specific gene. Our search is performed on a set of 2,758 unique human genes encoding cell-surface proteins from the human surfaceome database, including the CAR targets for solid tumors in clinical trials (**Fig. 1B**, **Methods**). Although published experimental work has explored up to three-gene ^19^ CAR circuits, we extended this by comprehensively screening N-gene circuits (N = 2, 3, 4, or 5) utilizing AND, OR, and NOT logic gates. Importantly, even when considering different genes as interchangeable within a symmetric circuit, a vast number of non-equivalent circuits exist. Specifically, the numbers of unique logic gate designs for two, three, four, and five input genes are 6, 20, 70, and 252, respectively (**Fig. 1A**, **Methods**). Because ‘!(A & B)’ is equivalent to ‘(!A) | (!B)’ and because ‘!(A | B)’ is equivalent to ‘(!A) & (!B)’, it follows that all NOT gates can take as input a (single gene) variable rather than a subexpression. Then, by treating a gene (G) and its negated form (!G) as distinct possible input variables, we eliminated the NOT logic gate, considering only AND and OR logic gates in the search. Eliminating NOT gates reduces the numbers of unique gate arrangements in a circuit to 2, 4, 12, and 40 for two, three, four, and five genes, respectively (**Methods**). However, even for simple two-gene logic gates, millions of possible circuits exist because of the large number of possible gene variables, rendering exhaustive searches infeasible on single computers for N ≥ 3.

**Figure 1.**
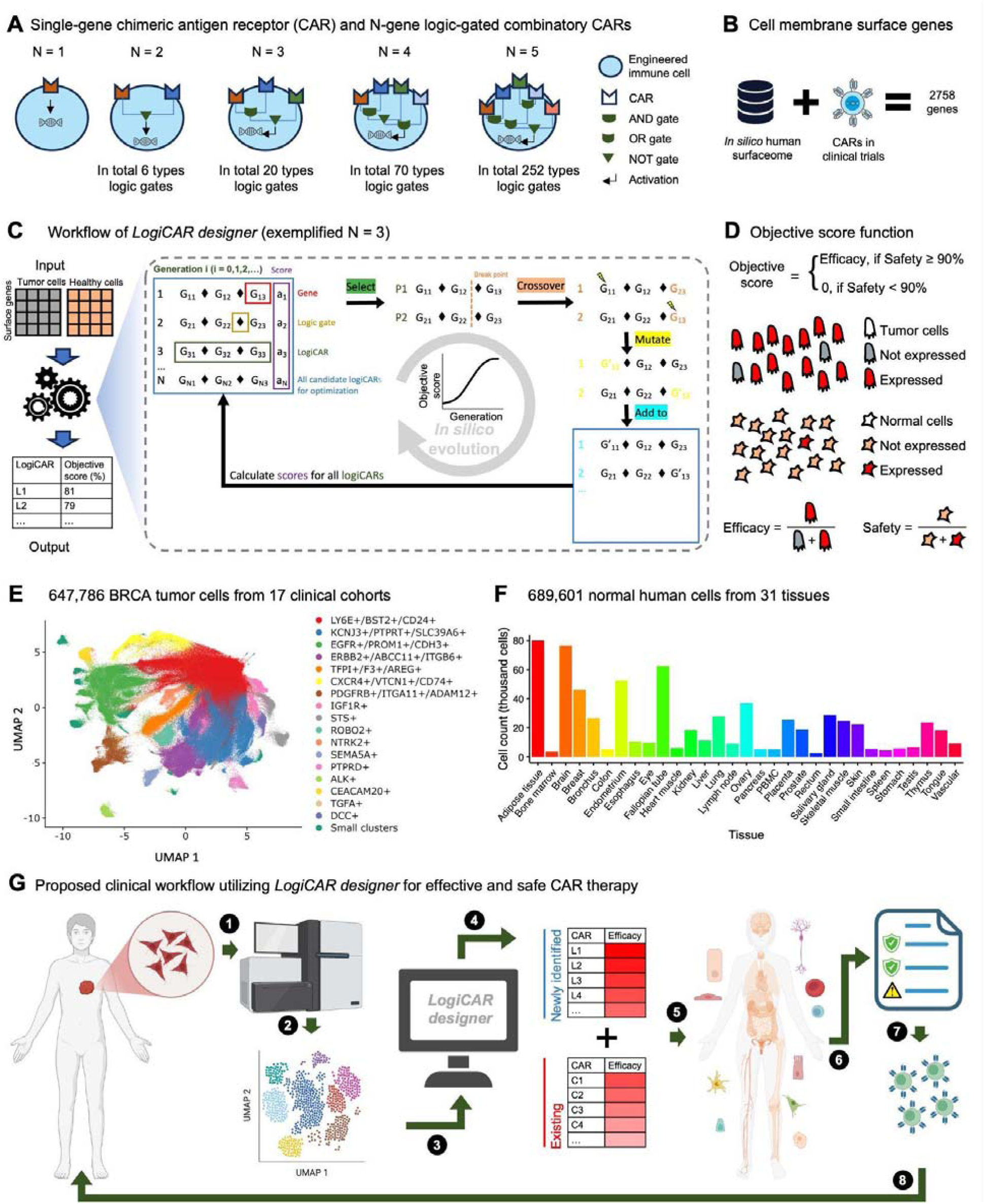
| Study Design Overview and Application. **A.** An illustration of single-gene chimeric antigen receptor (CAR) and N-gene CAR circuits. The number of genes, N, in a circuit is up to five in this study. The total number of designs for each different number of genes is shown. **B.** Curation of genes encoding cell membrane surface proteins. The list of eligible genes is derived from a union of the *in silico* human surfaceome and CAR targets in clinical trials of solid tumors (**Methods**). The total number of cell membrane surface genes is 2,758. **C.** Workflow of the *LogiCAR designer* algorithm, exemplified by N = 3 genes in a logic-gated combinatorial CAR. Left panel: input, algorithm schematic, and output. The input for *LogiCAR designer* consists of two single-cell gene expression matrices—one for tumor cells and one for healthy human tissue cells. The output is a list of identified top-ranked CARs based on the objective score function. Right panel: An illustration of the *in-silico* evolution mechanism of the *LogiCAR designer* algorithm. A set of candidate circuits was generated randomly from the gene pool (panel B) as Generation 1, and their objective scores were calculated. The top-scoring circuits were selected for gene exchange between circuits (Crossover) and gene replacement (Mutation). This new set of circuits becomes the next Generation entering the new evolution cycle. This process is repeated until the objective scores converge to a plateau. **D.** The objective score function. The objective score is defined as the proportion of tumor cells that express the CAR under test, with a preset safety threshold for the proportion of normal cells that does not express the CAR under test. In this study, the safety threshold was set at 90%. **E.** The breast tumor cells collected for this study from 15 public and 2 in-house cohorts. The cells were clustered based on the expression profiles of surfaceome genes. Clusters were named based on a few most differentially expressed surfaceome genes. Clusters with fewer than 5,000 cells were merged to form a mixed cluster labeled “Small clusters.” **F.** The 689,601 normal human tissue cells from 31 public datasets from the human Protein Atlas (HPA). **G.** Proposed clinical workflow for CAR therapy based on rational design. The process begins with the dissociation of tumor tissue from a cancer patient into single cells, which are then prepared for single-cell gene expression analysis. Following sequencing, the cells are annotated to distinguish tumor cells from non-tumor cells. The expression matrix of tumor cells is then input into the *LogiCAR designer* tool to identify potential CAR circuits. The efficacy of these newly identified circuits is evaluated alongside existing CARs using computational models. Next, the candidate CARs undergo safety evaluations across major human tissues and cell types, assessed at both RNA and protein expression levels. A comprehensive efficacy and safety profile report for the potential CARs is then generated in detail. Based on multiple considerations, including time, financial resources, and disease severity, clinicians select a single-or a multi-gene CAR circuit. CAR-engineered immune cells are then manufactured and injected into the patient.

To overcome the challenge of searching in such an exhaustive combinatorial space, we developed *LogiCAR designer* (**Fig. 1C**) to identify CAR circuits. *LogiCAR designer* takes two single-cell gene expression matrices as input: one from tumor cells and one from healthy human tissues, which is the same for all patient sets of the same cancer type. The tumor cell input varies by data set and can be from multiple cohorts, a single cohort with multiple patients, or a single patient. For assessing safety, we built a single-cell RNA sequencing (scRNA-seq) gene expression matrix from 31 healthy human tissues across 31 datasets as the healthy tissue input. *LogiCAR designer* outputs ranked lists of N-gene CAR circuits (N = 1–5) based on an objective score (**Fig. 1C**, left panel).

*LogiCAR designer* uses an iterative genetic algorithm ^20^ (**Fig. 1C**, right panel). For a given N (e.g., N = 3) and target circuit type (e.g., ‘G1 | G2 | G3’) as “genotype”, an initial random population of circuits is generated by randomly selecting an ordered list of genes from the cell surfaceome and assigning them to be the input genes of the circuit. The *objective score* of each candidate circuit is then calculated as its “fitness” (to determine the likelihood that it will be retained in further search iterations of the algorithm). The objective score estimates the efficacy of tumor cell targeting while satisfying a predefined safety threshold. *Tumor cell targeting efficacy* is defined as the *percentage of tumor cells* expressing a combination of inputs (i.e., that evaluate to 1 in the circuit). The *safety threshold* was set at 90% *sparing of input healthy single cells* (i.e., evaluating to 0 in the circuit; **Fig. 1D**), consistent with prior work ^11,17^. The search / optimization process then continues iteratively across successive generations until convergence is obtained. That is, in each iteration (generation), the circuits with high objective scores are selected for the next generation with probabilities proportional to their scores. Selected “parental” circuits undergo crossover (gene exchange) to produce “offspring.” To introduce variability, genes in the offspring can be “mutated” to random genes from the cell surfaceome with a certain probability (**Methods**). This selection-crossover-mutation process iterates until a new generation replaces the previous one, repeating until a stopping criterion is met when the scores of the solutions found cease to improve (**Methods**). We focused on BRCA, the most common female cancer in the United States (U.S.). To ensure unbiased screening and assess generalizability, we curated a comprehensive BRCA dataset comprising scRNA-seq and single-nucleus RNA sequencing (snRNA-seq) data. This dataset comprises 1,739,931 cells, including 622,945 tumor cells and 1,116,986 non-tumor cells, derived from 15 publicly available cohorts and two in-house cohorts (**Fig. 1E; Supplementary Fig. 1**; **Table 1**). These data collectively represent 342 patient samples (**Table 1**, **Supplementary Table 1**). It spans all major BRCA subtypes, primary and metastatic tumors, and diverse sequencing platforms (**Table 1**). As expected, the tumor cells exhibited substantial heterogeneity, clustering based on surfaceome expression (**Fig. 1E**). For the safety score calculation, we used the scRNA-seq expression matrix of 689,601 normal human tissue cells from 31 public datasets provided by the Human Protein Atlas (HPA) project (**Fig. 1F**).

**Table 1.**
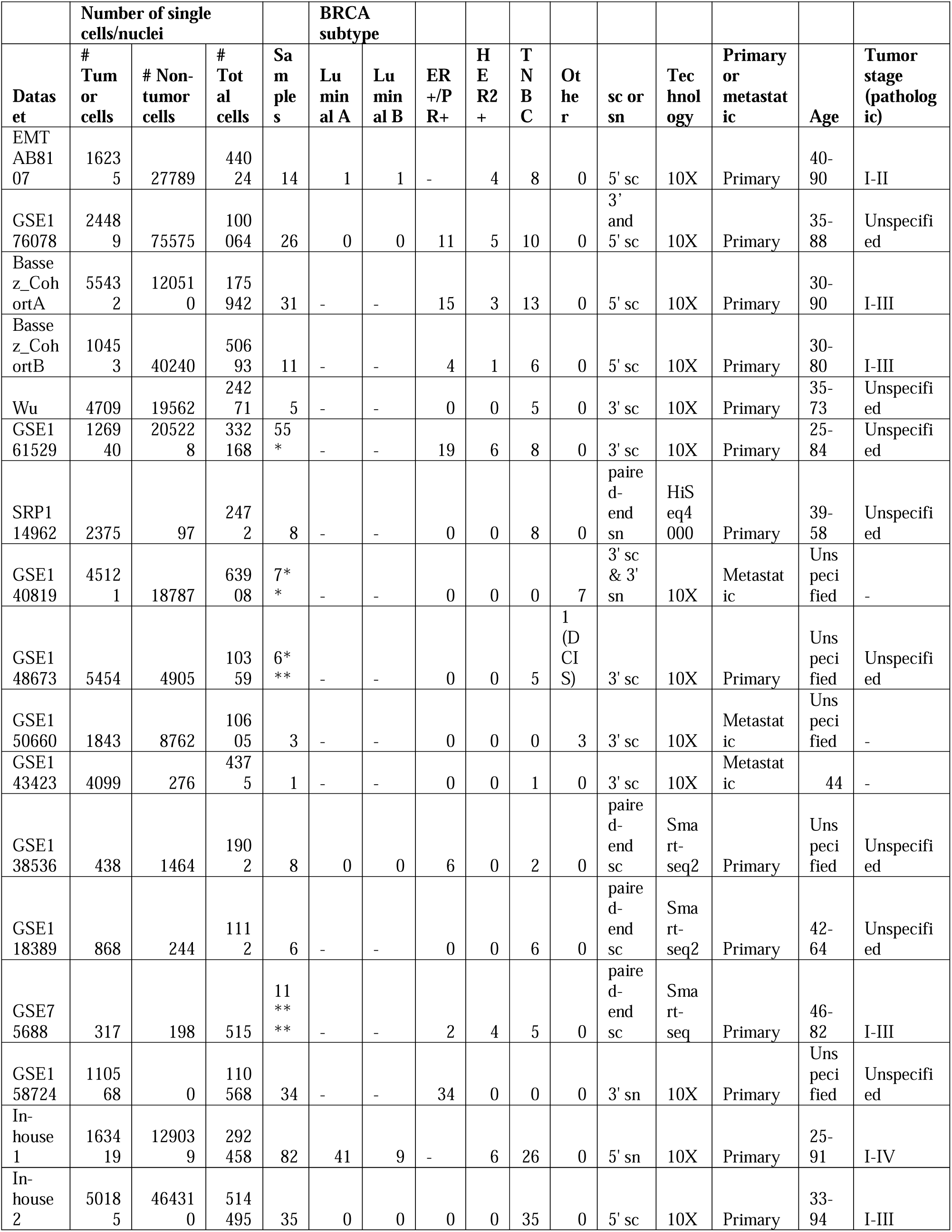
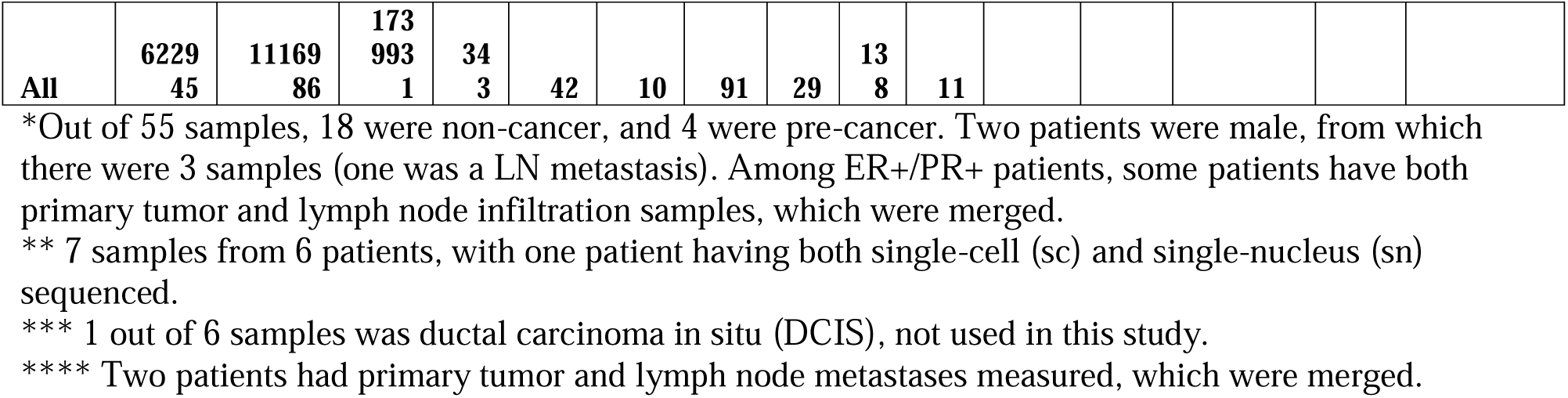
Sequencing and clinical information of the 17 BRCA cohorts in this study.

In a potential clinical workflow for CAR therapy, we foresee two main applications for the *LogiCAR designer* approach: (1) In the first, we applied it to analyze the breadth of the cohorts described above to identify and estimate the quality of the highest scoring logic CAR circuits that are shared across many patients in as many as possible datasets. Those could serve as promising *shared candidate logic circuits* for the design and testing of the next generation of CARs. But second, (2), in a future precision oncology scenario, with the development of fast ways to engineer tailor-made CARs, a practitioner could generate single-cell data (either the transcriptomic data or proteomics data) from a patient or cohort and deploy *LogiCAR designer* to select promising new *individualized candidate logic circuits*. The efficacy of those, as we shall show, usually markedly surpasses that of shared gates. Crucially, a comprehensive safety profile, including RNA and protein expression in different normal tissues and cells, is evaluated for all candidate gate designs. Based on the efficacy and safety report of candidate CAR logic circuits, the clinicians can then make more informed decisions on which CAR circuit to use, including practical considerations of time, financial resources, and disease severity. (**Fig. 1G**).

### Designing more effective CAR circuits for breast cancer that are shared across many cohorts and patients

We begin by establishing the need for developing an optimization algorithm to solve the research question in hand, and then set out to systematically compare the estimated running times of *LogiCAR designer* versus exhaustive brute force search by measuring those for an increasing number of targets, N, in the desired circuit. Setting *LogiCAR designer*’s time complexity at N = 2 as the base value of 1, we find that the exhaustive search time complexity at N = 2, 3, 4, and 5 grows exponentially, with magnitudes of 1.1 × 10^1^, 3.3 × 10^4^, 7.9 × 10^7^, 1.5 × 10^11^, respectively, while *LogiCAR designer*’s time complexity only grows linearly with the input size, 1, 10, 48, and 200 (**Fig. 2A**; see **Methods**). For example, running a three-gene design with *LogiCAR designer* takes ∼10 minutes on an M3 Macbook with 36 GB RAM and 12 CPU cores, compared to an estimation of ∼550 hours for the exhaustive search. *LogiCAR designer* demonstrates efficient convergence across varying tumor cell counts (500, ∼8,000, and ∼74,000) (**Supplementary Fig. 2A**), showing its scalability and independence from input size—a challenge for methods like integer linear programming (ILP) ^21^—while consistently identifying global optima in N = 2 cases, as validated by exhaustive search.

**Figure 2.**
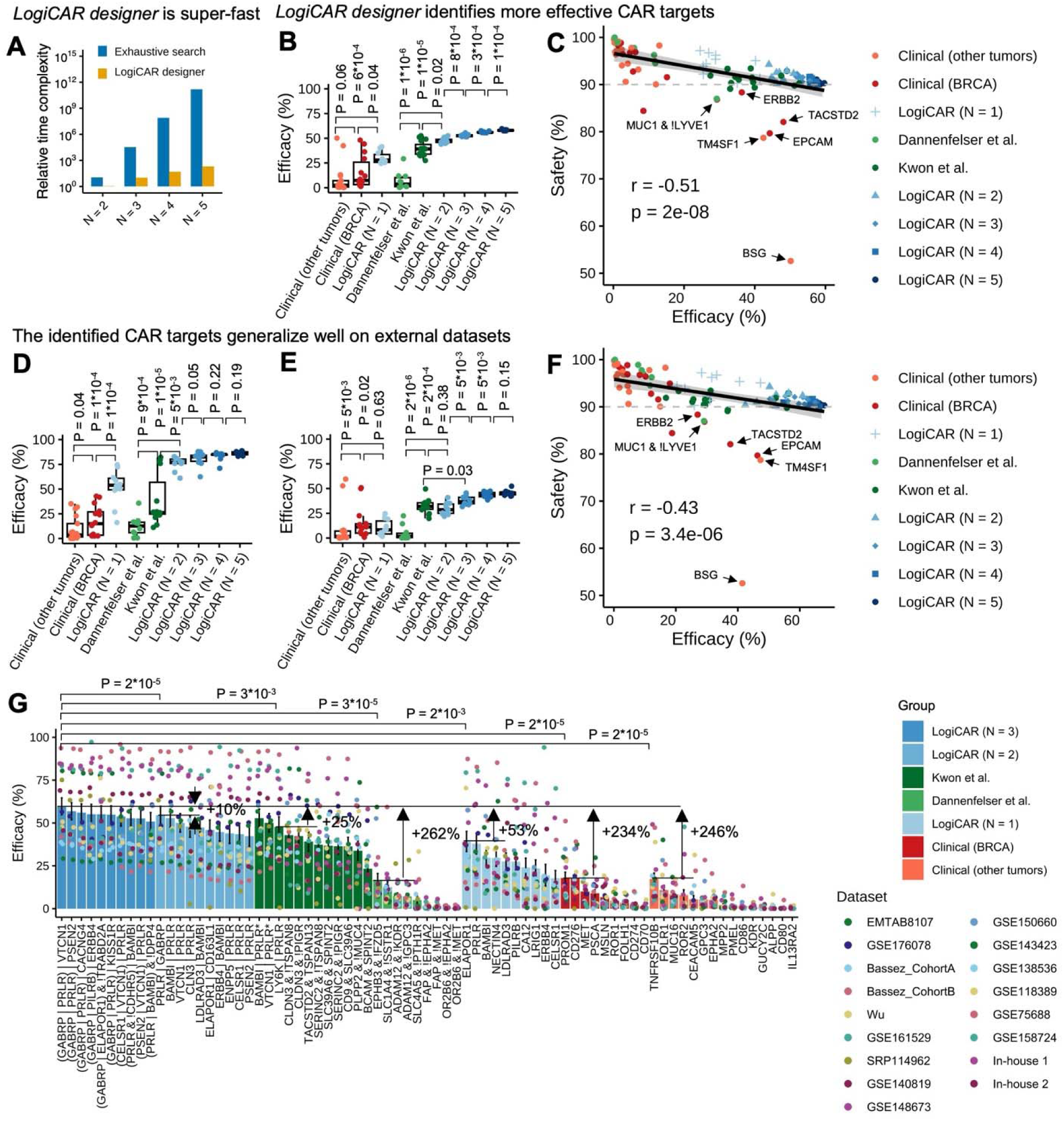
| The efficacy and safety landscape of top candidate shared (between patients) LogiCAR circuits in breast cancer. **A.** Comparison of relative running time (relative time complexity) between *LogiCAR designer* and exhaustive search for identifying optimal CAR circuits with gene numbers N = 2, 3, 4, 5. The time complexity of *LogiCAR designer* at N = 2 is set to 1 (see time complexity calculation in **Methods**). **B.** Comparison of efficacy between the top 10 newly identified target circuits for N = 5, 4, 3, 2, 1, respectively, and the previous computationally identified single- or multi-gene circuits, clinical trial single targets for BRCA, and clinical trial targets for other solid tumors. The Wilcoxon U-test was used, and two-tailed p-values are shown. Efficacy was calculated from the randomly sampled 74k tumor cells from the public cohorts. **C.** Safety versus efficacy for the CAR circuits in different groups in B. Safety was calculated from the randomly sampled 31k normal cells from the HPA curation. **D, E**. Same analyses as B, but evaluated in two in-house cohorts of ∼160k and ∼52k tumor nuclei / cells, respectively. **F**. Safety vs. efficacy averaged over the two in-house cohorts in panels D and E. **G**. Efficacy comparison of CAR circuits (N ≤ 3) with safety scores higher than 90%. Efficacy in all the 17 cohorts was calculated individually using all measured BRCA tumors. Error bars denote the standard error of the mean. If a circuit appears in multiple groups, its subsequent occurrences are marked with an asterisk.

We next turn to use *LogiCAR designer* to identify shared CAR circuits for BRCA, our main goal. To this end, as input for the search, we randomly sampled 500 tumor cells per patient from the 15 public discovery cohorts (totaling 72,433 cells, ∼18% of all measured tumor cells in discovery cohorts) and 1,000 cells per HPA normal tissue dataset used to determine safety (totaling 31,000 cells, ∼4% of all measured normal cells). The resulting maximal tumor targeting efficacy, *i.e.*, the estimated percentage of tumor cells targeted by the *LogiCAR designer*-selected circuit, for five, four, three, two, and one-gene CARs was a coverage of 60%, 58%, 56%, 53%, and 41% of all tumor cells respectively (**Fig. 2B**). Notably, the 2-gene LogiCAR circuits we identified demonstrated significantly higher efficacy compared to previously computationally identified 2-gene LogiCAR circuits derived from bulk RNA-seq (Dannenfelser et al.) or scRNA-seq (Kwon et al.) data (p = 1 × 10^-5^ and 0.02, respectively; **Fig. 2B**) ^12,16^. Similarly, the single-gene CAR targets we identified exhibited significantly greater efficacy compared to single-gene targets under clinical trial investigation, both for BRCA and other solid tumors (p = 0.04 and 6 × 10^-4^, respectively; **Fig. 2B**). Importantly, all our identified circuits, in difference from several existing ones, surpassed the safety threshold (**Fig. 2C**; **Supplementary Fig. 2B**). Generally, safety and efficacy were negatively correlated (**Fig. 2C**), as one would expect. Of note, some previously computationally identified circuits (e.g., ‘*MUC1 & !LYVE1*’, ‘*EPCAM* & !*SSTR1*’, ‘*CD24 & TM4SF1*’) and clinical trial targets (e.g., *BSG*, *EPCAM*, *TACSTD2*, *TM4SF1*, *ERBB2*) fell below the safety threshold.

To test the generalizability of these findings, we evaluated target efficacy in two test cohorts: a large in-house snRNA-seq cohort (82 patients) across all BRCA subtypes (41 luminal A, 9 luminal B, 6 HER2+, and 26 TNBC), and a scRNA-seq cohort of 35 TNBC patients recently published ^22^ after our analysis started (**Methods**). The newly identified LogiCAR circuits consistently maintained their high scores and significantly outperformed previous and clinical trial targets also in these completely independent external validation cohorts (**Fig. 2D**, **E**). Interestingly, computational circuits derived from bulk RNA-seq data performed poorly across both the discovery and validation datasets (**Fig. 2B**, **D**, **E**), likely reflecting their inability to address tumor heterogeneity. While circuits identified from prior scRNA-seq analyses performed comparably to our two-gene LogiCAR circuits in the TNBC cohort (**Fig. 2E**), they performed significantly worse in the in-house snRNA-seq cohort (p = 5 × 10^-3^; **Fig. 2D**), suggesting possible overfitting of their models to scRNA-seq data. We also observed a similar negative correlation between safety and efficacy in the test cohorts (**Fig. 2F**), reinforcing the trends observed in the discovery cohorts (**Fig. 2C**). Finally, we noted diminishing returns in efficacy improvements as the number of genes increased beyond three (N = 3 to N = 4 or 5), with gains often becoming statistically insignificant (**Fig. 2D, E**). Given the experimental challenges associated with designing and implementing multi-gene CAR circuits ^23^, we focused on circuits with N ≤ 3 in the following in-depth analysis.

We comprehensively evaluated and ranked the efficacy of newly identified (N ≤ 3) and existing CAR circuits (with safety score > 90%) using all 622,945 tumor cells across 17 individual cohorts (note that only 72,433 tumor cells from 15 public cohorts were used for identifying the optimal CAR circuits). As evident in **Fig. 2G**, the best identified circuit, ‘*GABRP | PRLR | VTCN1*’, achieved 60% efficacy, which was 234% and 246% more effective than the best targets in clinical trials for BRCA or other solid indications, respectively (**Fig. 2G**). Interestingly, the top two circuits identified by previous computational methods using scRNA-seq, ‘*BAMBI | PRLR*’ and ‘*VTCN1 | PRLR*,’ overlapped with our identified two-gene LogiCAR circuits, ranking 2nd and 3rd, respectively (**Fig. 2G**). This overlap occurred despite the use of different computational approaches and datasets, underscoring the robustness of these circuits (see **Methods** and **Supplementary Table 2** for complete lists of previous computationally identified circuits and targets in clinical trials).

### Characterizing the safety landscape of identified shared CAR circuits

Beyond efficacy, safety is a paramount consideration for CAR T-cell therapy ^1,11^. To develop as comprehensive a safety evaluation as possible, we analyzed the expression patterns of potential CAR circuits in major human tissues and cell types, classifying tissues and cell types from scRNA-seq and protein immunohistochemistry (IHC) data into 18 and 19 major categories, respectively (**Fig. 3A**; **Methods**). For the expression of combinations, an “equivalent expression” to single genes or proteins is derived based on the logic gates. For example, the “equivalent expression” of G1 & G2 is calculated as the minimum of G1 and G2. Detailed calculations for the “equivalent expressions” of other logic gates are provided in the **Methods** section.

**Figure 3.**
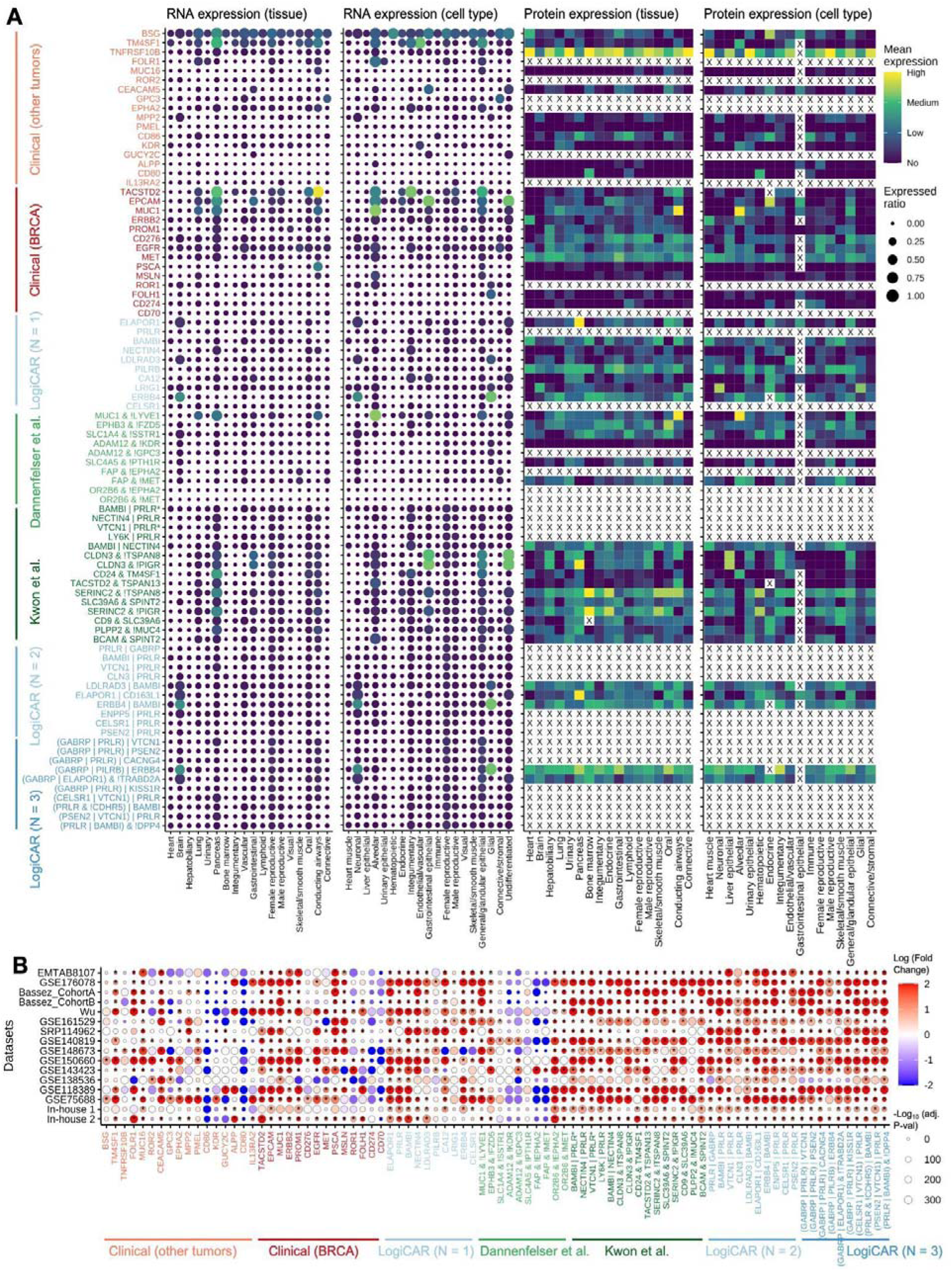
| Characterizing the Safety Landscape of leading shared CAR circuits. **A.** RNA (left two panels) and protein (right two panels) expression profiles of potential CAR circuits across major human tissues and cell types. “X” denotes missing data. The color gradient shows the relative expression level, and the size of the circle indicates the ratio of circuit-positive cells in all cells. The “High,” “Medium,” “Low,” and “No” expression levels for proteins are based on the original HPA IHC annotations. To quantify RNA expression levels, we applied a log1p transformation: raw gene expression counts for each cell were normalized to 10,000 counts per cell and transformed using the natural logarithm of the normalized counts plus one. Expression levels in single cells or nuclei were then categorized in the plot as follows: values greater than 3 were capped at 3 and labeled as “High,” values of 2 as “Medium,” values of 1 as “Low,” and values of 0 as “No” expression. If a circuit appears in multiple groups, its subsequent occurrences are marked with an asterisk. **B.** RNA-level safety landscape of LogiCAR-designed circuits and single-gene CARs in the tumor microenvironment (TME) across BRCA datasets. The heatmap depicts the fold changes of log-normalized single-cell RNA expression between tumor cells and non-tumor cells for each CAR circuit. Statistical significance was determined using a two-tailed Wilcoxon test, with p-values adjusted for multiple comparisons using the Bonferroni method. Data points with an adjusted p-value < 0.05 and a fold change > 2 are marked with an asterisk (*). The dataset GSE158724 was excluded from this analysis due to the absence of non-tumor cells measured. See more detailed log-normalized gene expression profile in tumor and non-tumor cells in **Supplementary Fig. 4**.

At the RNA level, we found that existing circuits could be frequently expressed in normal tissues and cell types (**Fig. 3A**, left two panels). For example, a target in clinical trials, *BSG*, was quite frequently expressed in multiple normal tissues and cell types. *MUC1* and *MUC1*-containing circuits were expressed in lung and pancreas tissues and alveolar and general/glandular epithelial cells. *EPCAM* and *EPCAM*-containing circuits were expressed in pancreas and gastrointestinal tissues and alveolar, gastrointestinal epithelial, general/glandular epithelial, and undifferentiated cells. *TACSTD2* (encodes TROP2) showed moderate expression in pancreas, low to moderate expression in oral tissue, and high expression in conducting airway tissue, and low to moderate expression in alveolar, integumentary, and general/glandular epithelial cells. *TM4SF1* was expressed in pancreas and conducting airway tissues and alveolar, endothelial/vascular, and general/glandular epithelial cells. The newly identified *ERBB4* and *ERBB4*-containing circuits show low to moderate expression in brain tissue and neuronal and glial cells (**Fig. 3A**, left two panels).

At the protein level, the target in clinical trials *TNFRSF10B* (clinical trials NCT03638206, NCT06251544) showed high expression in multiple tissues and cell types. *MUC1*^24^ (NCT04020575) and *MUC1*-containing circuits showed high expression in conducting airway tissue and alveolar cells. *EPCAM* and *BSG* showed moderate to high expression in endocrine cells. *ERBB2* (HER2) showed moderate expression in heart tissue and heart muscle cells and low to moderate expression in alveolar cells (**Fig. 3A**, the two right panels). While the newly identified *ELAPOR1* and ‘*ELAPOR1* | *CD163L1*’ (**Fig. 2G**) showed high expression in pancreas, this can be addressed by combining with *TRABD2A* using NOT in the circuits ‘(*GABRP* | *ELAPOR1*) & !*TRABD2A*.’ Due to the substitution of *CD163L1* with *GABRP*, the efficacy also increased (**Fig. 2G**). This is an example that rationally combining multiple genes in a CAR can simultaneously enhance efficacy and safety. Some single-gene and multiple-gene CAR circuits lacked protein data in the HPA database, which is denoted as “X” in the right two panels of **Fig. 3A**.

We summarized the safety and efficacy profiles of each CAR circuit, showing the maximum mean expression across all tissue or cell types at the RNA and protein levels, respectively (**Supplementary Fig. 3**). Notably, the newly identified CAR circuits generally had similar or better safety profiles compared to the existing CAR circuits. In particular, ‘(*GABRP* | *ELAPOR1*) & !*TRABD2A’* had high efficacy and relatively low expression across tissues and cell types at both RNA and protein levels (**Supplementary Fig. 3**). Many *PRLR*-containing circuits lack protein level data and need further investigation of safety. However, multiple *SERINC2-*, *CLDN3-*, and *MUC1*-containing previous computational circuits, and some clinical trial targets like *TACSTD2* (NCT05969041), *BSG* (NCT03993743), *EPCAM* (NCT02915445), *TM4SF1* (NCT05673434, NCT04151186), and *TNFRSF10B* (NCT03638206) require careful safety evaluation due to their moderate to high expression in some normal tissues or cells (**Fig. 3A**; **Supplementary Fig. 3**).

To further evaluate the tumor-specificity of CAR circuits, we investigated their expression patterns within the tumor microenvironment (TME), comparing tumor cells to non-tumor cells in patient tumor samples. An ideal CAR circuit for therapeutic applications should exhibit significantly higher expression in the former compared to the latter, ensuring both efficacy and safety. Our analysis revealed that many single-gene CAR targets in clinical trials, as well as CAR circuits identified through previous bulk RNA-seq studies, demonstrated poor discriminatory ability between tumor and non-tumor cells in the TME (**Fig. 3B**; **Supplementary Fig. 4**). These findings underscore the limitations of bulk RNA-seq data in resolving tumor-specific signals.

Strikingly, several novel single-gene CAR targets identified in this study, such as *ELAPOR1* and *BAMBI*, consistently exhibited a fold-change > 2 in tumor versus non-tumor cells across all BRCA datasets, with adjusted p-values < 0.05 (**Fig. 3B**; **Supplementary Fig. 4**). Furthermore, multiple CAR circuits incorporating these targets—including ‘*BAMBI* | *PRLR*, *BAMBI* | *LDLRAD3’*, ‘*BAMBI* | *ERBB4’*, ‘*ELAPOR1* | *CD163L1’*, *‘*(*PRLR* & !*CDHR5*) *| BAMBI’*, and *‘*(*PRLR | BAMBI*) & !*DPP4’*—also achieved a significant and consistent fold-change > 2 in tumor versus non-tumor cells across all BRCA datasets (**Fig. 3B**; **Supplementary Fig. 4**). In contrast, none of the existing CAR circuits derived from clinical trials or prior computational studies demonstrated comparable tumor-specific discriminatory power (**Fig. 3B**; **Supplementary Fig. 4**).

These results are remarkable considering that our CAR circuit identification pipeline did not explicitly incorporate non-tumor cells from the TME. The tumor-specific expression of these newly identified CAR targets and circuits highlights their potential as effective and safe therapeutic candidates.

### Patient-level assessment of the efficacy of leading shared CAR circuits

To provide a more clinically relevant assessment of the CAR designs we have identified, we evaluated the *patient-level* tumor-targeting efficacy and coverage of the candidate CAR circuits. We focused on 206 samples that had more than 500 tumor cells measured, out of a total of 342 patient samples from the 17 BRCA cohorts. We found that the newly identified individualized LogiCAR circuits have substantially stronger tumor-targeting efficacy at the patient level compared to previously identified computational circuits and those in clinical trials (**Fig. 4A**).

**Figure 4.**
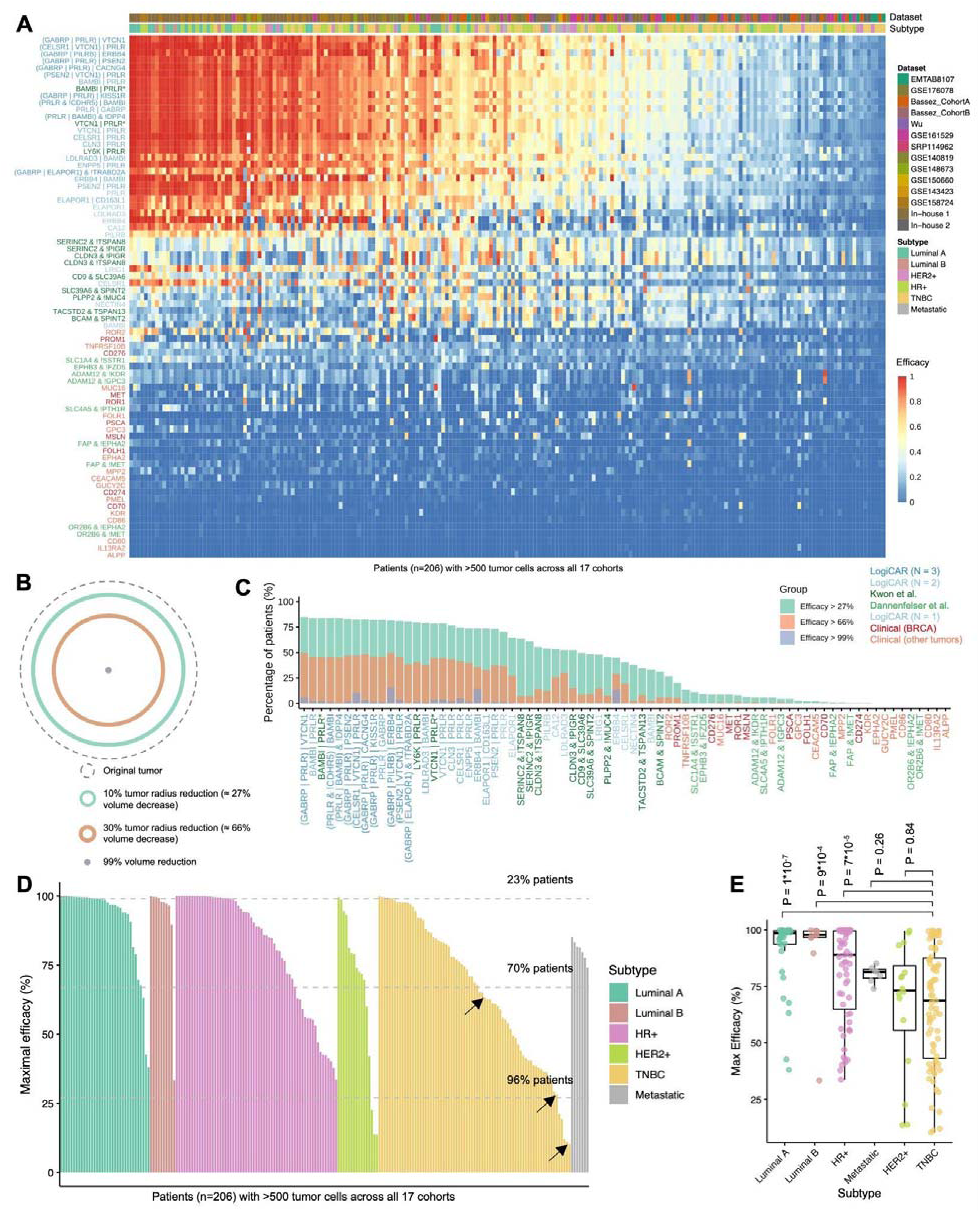
| Patient-level tumor-targeting efficacy of top candidate shared LogiCAR circuit designs in RECIST terms. **A.** Efficacy comparison of CAR circuits (N ≤ 3) with safety scores >90% across 206 patients (>500 tumor cells each) from 17 cohorts. Dataset origin and BRCA subtype are indicated for each patient. If a circuit appears in multiple groups, its subsequent occurrences are marked with an asterisk. **B.** Relationship between tumor volume reduction and radius reduction, assuming the tumor is a solid sphere. In clinical practice, a 10% reduction in tumor radius is considered a minimal pathological response, a 30% reduction is classified as a partial response, and the absence of visible tumors is defined as a complete response. To account for numeric precision, we use a >99% volume reduction as the threshold for a complete response. **C.** Efficacy distribution of potential CAR circuits across patients. For each CAR circuit, the percentages of patients achieving tumor-targeting efficacy thresholds of >27% (minimal pathological response), >66% (RECIST partial response), and >99% (complete response) are shown. If a circuit appears in multiple groups, its subsequent occurrences are marked with an asterisk. **D.** Maximum efficacy achievable for each patient using the optimal CAR circuit among the potential candidates. By optimizing CAR circuit selection for each patient, 23%, 70%, and 96% of patients achieve tumor-targeting efficacy thresholds of >99% (complete response), >66% (partial response), and >27% (minimal pathological response), respectively (thresholds indicated by dashed lines). However, 4% of patients exhibit tumor-targeting efficacy below 27% regardless of the CAR circuit used. The three arrows highlight three representative TNBC patients who will be examined in detail in **Fig. 5A, B, C**. **E.** Maximal efficacy in patients from different subtypes. P values are from a two-tailed Wilcoxon test.

We further analyzed the efficacy profiles of each CAR circuit. To this aim, we note that in clinical practice, a reduction of at least 10% but less than 30% in tumor radius could be considered a minimal pathological response ^25^, a reduction of at least 30% corresponds to a partial response based on the RECIST criteria ^26^, and the absence of visible tumors is regarded as a complete response. Assuming a tumor is a solid sphere, 10% or 30% radius reductions correspond to 27% or 66% reductions in tumor volume, respectively (**Fig. 4B**). To account for numerical precision, we set a >99% reduction in tumor volume as the threshold for a complete response. For each CAR circuit, we determined the proportion of patients whose tumor-targeting efficacy surpassed 27%, 66%, and 99%, as approximations of the different levels of clinical response (**Fig. 4C**).

Encouragingly, the newly identified LogiCAR circuits demonstrated substantial efficacy. For example, the LogiCAR circuit ‘*GABRP* | *PRLR* | *VTCN1*’ achieved the highest proportion of patients (85%) with tumor-targeting efficacy greater than the minimal pathological response 27% threshold. Similarly, both ‘*GABRP* | *PILRB* | *ERBB4*’ and ‘*GABRP* | *PRLR* | *VTCN1*’ showed the highest proportion of patients (50%) with efficacy greater than the 66% partial response threshold. For tumor-targeting efficacy that signifies complete response (greater than 99%), the LogiCAR ‘*GABRP* | *PILRB* | *ERBB4*’ outperformed all other circuits, with 16% of patients achieving this threshold (**Fig. 4C**).

As illustrated in **Fig. 4A**, tumor cells from individual patients exhibited varying responses to different LogiCAR circuits, emphasizing the potential benefits of designing patient-specific CAR therapies. To explore this further, we analyzed the theoretical maximum tumor-targeting efficacy achievable for each patient by matching the most effective LogiCAR circuits from our pool (**Fig. 4D**). Our analysis revealed that optimal CAR circuit-patient matches could achieve tumor-targeting efficacy exceeding 99% (indicative of a complete response) for 23% of patients. Additionally, 70% of patients could achieve efficacy greater than the 66% partial response threshold, while 96% of patients could reach efficacy levels exceeding 27%, indicative of a minimal pathological response.

However, 8 out of the 206 patients are likely to be complete non-responders to any of the CAR designs in our optimal set because they did not have any circuits with estimated tumor-targeting efficacy above 27% (**Fig. 4D**). Notably, the majority (5 out of 8) of these patients were TNBC cases, highlighting the higher heterogeneity and greater therapeutic challenges associated with this aggressive subtype. Consistently, when evaluating the best tumor-targeting efficacy achieved through optimal CAR circuit-patient matching across different subtypes, TNBC patients exhibited significantly lower efficacy compared to luminal A, luminal B, and HR+ (ER- or PR-positive) patients (p = 8 × 10^-8^, 8 × 10^-5^, 4 × 10^-5^, respectively; **Fig. 4E**). These findings underscore the importance of individualized CAR circuit design to enhance tumor-targeting efficacy, particularly for patients with heterogeneous or refractory tumor profiles, which we will address in the next section.

### CAR circuit selection for individual TNBC patients puts forward highly effective and safe ***patient-specific* tailored CAR circuits**

Tumor heterogeneity, both inter- and intra-patient, is a major challenge in solid tumors ^23^. Our findings revealed that the best-identified shared CAR circuit designs achieved an overall efficacy of ∼60% when evaluated across all 17 cohorts (**Fig. 2G**); while this is a major improvement over the efficacy and safety scores of extant solutions, it is still a far cry from the ambitions and goals of precise, personalized oncology. In addition, approximately 4% of patients had tumor-targeting efficacy scores below the threshold of minimal response, regardless of the LogiCAR circuit used (**Fig. 4D**). To address these limitations, we explored the potential of *individualized CAR circuit design,* which aims to leverage single-cell transcriptomics data to tailor the best possible LogiCAR circuits to individual patients.

We begin by reviewing the results achieved on three ‘case study’ TNBC patients from distinct cohorts, which were each selected due to being extreme by a different measure described below, to illustrate the benefits of this approach. The first is Patient 0135 (cohort GSE161529). This patient’s tumor, with 14,376 tumor cells, has the largest TNBC tumor cell population in our dataset. Without individualized CAR circuit design, the best tumor-targeting efficacy for this patient was 63%, nearing the threshold for a ‘partial response’ (**Fig. 4D**, left-most arrow). However, applying the *LogiCAR designer* to this patient’s tumor cells identified the individualized circuit, ‘(!*ITM2B | TACSTD2*) *& EMP2*,’ which achieved an efficacy of 89%, deep in the partial response zone. For perspective, the efficacy scores of existing clinical targets for BRCA or other solid indications are 15% and 20%, respectively (**Fig. 5A**, lower panel). The personalize optimized logic circuits effectively address tumor heterogeneity. For example, clinical trial targets like *FOLR1* (NCT03585764), *PROM1* (NCT03356782), and *CD276* (encodes B7H3; NCT04691713) were only expressed in distinct, small tumor cell subpopulations (**Supplementary Fig. 5**). Single-gene targets like *NECTIN4,* an emerging target in clinical study (NCT03932565, NCT06724835), had also moderate tumor-targeting efficacy in this patient (**Supplementary Fig. 5**). In addition, while some single-gene targets like *TACSTD2* and *EMP2* were widely expressed across tumor cells, they did not pass the safety threshold (**Fig. 2F**). Combining them with other genes such as *ITM2B* improved safety while maintaining efficacy, e.g. using the ‘(!*ITM2B* | *TACSTD2*) & *EMP2*’ circuit (**Supplementary Fig. 5**).

**Figure 5.**
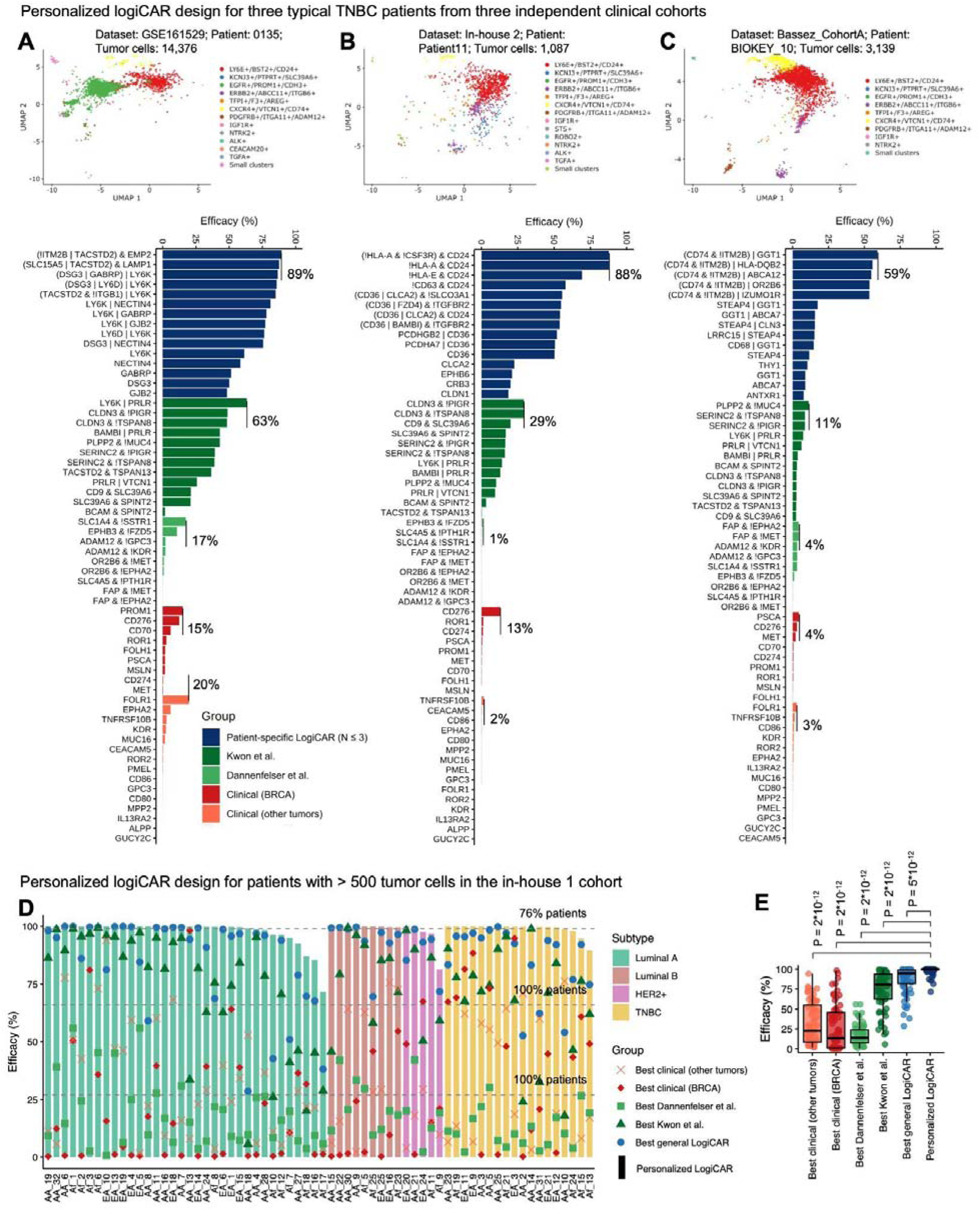
| Personalized CAR Circuit Design for TNBC Patients Overcome Inter-tumor Heterogeneity. **A.** Identification of individualized single-gene CAR targets and circuits for a TNBC patient from cohort GSE161529. Upper: UMAP plot of all 14,376 tumor cells from this patient. UMAP coordinates and clusters are the same as that in **Fig. 1E**. Lower: Comparison of the efficacy of potential CAR circuits for this patient. The potential CAR circuits include the top 5 best patient-specific N-gene circuits identified by running *LogiCAR designer* on this patient for each N = 1, 2, 3; the previously computationally identified circuits with safety scores higher than 90%; and the clinical trial targets with safety scores higher than 90%. If a circuit appears in multiple groups, its subsequent occurrences are marked with an asterisk. **B.** Same as Panel A, except the TNBC patient was from in-house cohort 2. **C.** Same as Panel A, except the TNBC patient was from Bassez CohortA. **D.** Tumor-targeting efficacy of different types of optimal CAR circuits in 66 individual patients with more than 500 tumor nuclei measured in the in-house cohort 1. Patients were grouped by subtype and ranked by best efficacy using personalized LogiCAR circuits. By optimizing CAR circuit design for each patient, 76%, 100%, and 100% of patients achieved tumor-targeting efficacy thresholds of >99% (complete response), >66% (partial response), and >27% (minimal pathological response), respectively (thresholds indicated by dashed lines). **E.** Comparison of the maximal efficacy achieved in 66 patients from the in-house cohort 1 using different groups of CAR circuits. P-values were calculated using a two-tailed Wilcoxon test.

The second patient is Patient 11 (in-house cohort 2). Patient 11, with 1,087 tumor cells, exhibited a best tumor-targeting efficacy of 29% without individualized design, corresponding to a ‘minimal pathological response’ (**Fig. 4D**, middle arrow). The individualized LogiCAR circuit, *‘*(!*HLA-A* & !*CSF3R*) & *CD24*,’ achieved an efficacy of 88%, dramatically outperforming the best targets identified by previous computational methods (29% and 1%, respectively) and clinical targets for BRCA or other solid indications (13% and 2%, respectively) (**Fig. 5B**, lower panel).

The third patient is Patient BIOKEY_10 (cohort Bassez_CohortA). This patient had the lowest shared tumor-targeting efficacy among all 206 patients without using individualized CAR circuit design, rendering the tumor entirely unresponsive to any shared CAR logic circuits (**Fig. 4D**, rightmost arrow; **Fig. 5C**). The individualized LogiCAR circuit, *‘*(*CD74* & !*ITM2B*) *| GGT1*,’ however, achieved an efficacy of 59%, approaching the partial response threshold of 66% (**Fig. 5C**, lower panel).

To systematically evaluate the potential of personalized CAR circuit design, we applied *LogiCAR designer* to in-house 1 cohort, which encompasses all major BRCA subtypes (**Fig. 5D**). Among the 66 patients with >500 tumor nuclei, the best shared CAR circuits achieved the following mean efficacies: *ROR2* and *PROM1*, targets currently under clinical investigation, achieved efficacy scores of 32% and 26%, respectively; the bulk RNA-seq-derived circuit *’SLC1A4* & (!*SSTR1*)’ had an efficacy score of 17%; the scRNA-seq-derived circuit *’LY6K* | *PRLR*’ had an efficacy score of 73%; and the LogiCAR designer-derived circuit ’(*GABRP* | *PILRB*) | *ERBB4*’ achieved an efficacy score of 87%.

Remarkably, personalized CAR circuit designs resulted in tumor-targeting scores equivalent to complete response levels in 76% of patients, and at least partial response levels in 100% of patients (**Fig. 5D**). Quite strikingly, the mean efficacy score of the individually tailored top CAR circuits across all patients was 98% (median: 99.9%), markedly surpassing all shared CAR circuits (**Fig. 5E**). Detailed results, including the top-ranked individualized CAR circuits and their efficacy, are provided in **Supplementary Table 3**.

## Discussion

In recent years, CAR T-cell therapies have been proven efficacious for B cell-lineage malignancies ^27,28^. However, the development of CAR T cell therapy for solid tumors has not yet led to any FDA approvals ^1^. One of the main reasons for the slower progress in the latter is the challenge of balancing efficacy and safety. For example, circuits expressed in a higher percentage of cancer cells are more likely to be expressed in non-malignant normal cells as well. This results in the observed negative correlation between the efficacy and safety of potential CAR circuits (**Fig. 2** **C, F**). Our study is the first to systematically evaluate the potential of logic-gated multi-gene CAR design to resolve such issues, focusing on BRCA but laying the computational foundation for the design of CAR cell therapies for a broad spectrum of cancer types.

We curated a large BRCA dataset (∼ 2 million cells / nuclei with > 620k from tumor cells) covering the common BRCA subtypes, enabling comprehensive evaluation of tumor heterogeneity and target efficacy. We identified new single-gene (e.g., *ELAPOR1, PRLR, BAMBI, LDLRAD3*, *NECTIN4*) and multi-gene (e.g., ‘*GABRP* | *PRLR* | *VTCN1’*, ‘(*GABRP* | *ELAPOR1*) & !*TRABD2A’*, ‘*LDLRAD3* | *BAMBI’*) circuits, with the complete set of N-gene LogiCAR designs (N = 1, 2, 3, 4, 5) available in **Supplementary Table 4**. Reassuringly, several top targets identified in our logic circuits are already in preclinical studies (e.g., *PRLR* in antibody-drug conjugates (ADCs) ^29^, and *NECTIN4* in CAR therapies ^30^). Our comprehensive cross-tissue and cell-type safety evaluation of the top LogiCAR circuits provides crucial insights into their further possible clinical development. For instance, our observation of HER2 protein expression in cardiac muscle cells and alveolar cells aligns with documented lung, cardiovascular, ^31,32^ and renal toxicities ^33^ in clinical studies. By incorporating NOT gates (!G) into circuit designs, *LogiCAR designer* aims to mitigate potential toxicities from CAR T cells by identifying gene combinations that would prevent T cell activation upon interaction with normal cells expressing the target antigen.

Our genetic algorithm is computationally highly efficient. For example, one three-gene design with 10 random seeds takes 10–20 minutes on a typical laptop. Running all four three-gene designs on a typical laptop takes about an hour, compared to an estimate of >450 days for the exhaustive search (**Methods**). The algorithm consistently found the global optimum for N = 2 genes, and its convergence is relatively independent of input size (**Supplementary Fig. 2A**; **Methods**). *LogiCAR designer* can be implemented for single-cell transcriptomics-based CAR circuit identification on personal computers in clinic offices, making it a practical approach if clinical personalized oncology advances to a stage at which scRNA-seq data are collected to guide individualized patient treatment. In earlier single-cell studies, we have made the argument that collection of single-cell data is cost effective and what is needed is a positive feedback research process in which collection of more single-cell data with clinical outcomes drives the development of new methods, such as *LogiCAR designer*, which can then be used in turn to analyze newly collected clinical single-cell data ^17,34^.

Our first key finding is that, at least when evaluating the efficacy and safety scores employed in this paper, the efficacy and safety of BRCA CAR circuits in clinical trials leaves much to be desired. Second, we find that the best shared CAR circuit achieves only ∼60% tumor-targeting efficacy across all patients (**Fig. 2G**). This reinforces the importance of individualized patient-specific designs, hoping that such approaches may become more feasible in the foreseeable future, further motivating their design and implementation. In three individual TNBC patients we presented as illustrative cases, such individualized CAR circuits markedly increased the tumor targeting efficacy by 25% (from 63% to 89%), 59% (from 29% to 88%), and 48% (from 11% to 59%) (**Fig. 5A, B, C**). At the cohort level, individualized CAR circuit design achieved a remarkable mean efficacy of 98% (median: 99.9%), with 76% of patients having complete response equivalent tumor-targeting levels while maintaining good safety scores (**Fig. 5D, E**). These results highlight the transformative potential of personalized LogiCAR designs to overcome tumor heterogeneity and achieve unprecedented efficacy, particularly for patients predicted to respond poorly to shared CAR circuits.

Our study has a few limitations. First, obviously, it is a computational study; however, we believe that the top identified logic circuits are likely to serve as promising candidates for further experimental validations to confirm their efficacy and safety in preclinical and clinical settings. These top candidates have been carefully evaluated via objective straightforward efficacy and safety scoring measures, based on analyzing the largest single-cell tumor microenvironment and healthy tissue data available. Bearing this in mind, to facilitate such future validation *in vivo* studies, one may choose to focus the search on a set of genes that have one-to-one orthologs in mice. Second, tumor-targeting efficacy was estimated based on RNA expression data, as has been used in similar previously published studies ^6–9,16^. However, it is well known that RNA expression may not always correlate with protein expression ^35–37^, and for some genes, this discrepancy could impact the results. The availability of accurate, high throughput single-cell proteomics data in the future would enable more precise analyses and validation of these circuits; however, for now, such data are lacking at a large scale. Additionally, we did not distinguish between tumor-targeting and tumor-killing efficacy; this distinction, which would require modeling effects of the tumor microenvironment, is not made in any of the similar studies summarized in the **Introduction** either. Thirdly, this study focused specifically on genes encoding cell surface proteins, which are most relevant for CAR-based therapies. However, *LogiCAR designer* can also be extended to analyze other subsets of protein-coding genes, which, while less pertinent to CAR therapy, could be highly valuable for other therapeutic modalities such as TCR therapies or ADCs. Finally, while this study focused on breast cancer, the most common cancer type among women in the U.S., *LogiCAR designer* is broadly applicable and can be readily adapted to address similar challenges in other tumor types and diseases, including autoimmune conditions ^38,39^ and aging-driven phenotypes via senolytic therapy ^40^.

In conclusion, our study presents a powerful, fast, and efficient computational framework for designing safer and more effective logic-gated CAR therapies. By analyzing single-cell transcriptomics with a highly effective genetic search algorithm, we identified novel shared LogiCAR designs of depth three whose tumor-targeting efficacy surpasses current designs by far. However, we also learnt that such cohort-shared designs leave many patients without effective CAR treatments. Encouragingly, we find that searching for individual designs that are precisely tailored by analyzing each patient’s tumor separately can identify effective and safe LogiCAR circuits for most of these patients, underscoring the exciting promise and potential of such personalized CAR designs going forward.

## Methods

### Description of the curated BRCA Clinical Cohorts

Below is a brief description of the 17 BRCA clinical cohorts used in this study. We also documented the major information, including molecular and clinical information, in **Table 1** and **Supplementary Table 1**. Most patients are female, but there are a few cases of male patients (**Table 1**).

#### Public cohorts

Below is a brief description of the 15 public cohorts curated in this study. All patient identifiers and BRCA subtype annotations were retrieved from the downloaded data. We observed inconsistencies in BRCA subtyping across studies. For example, some studies used receptor status-based classification (e.g., HR+ [ER+/PR+], HER2+, TNBC), while others used molecular status-based intrinsic subtype classification (e.g., luminal A, luminal B, HER2-enriched, basal-like, normal-like) or histological subtypes (e.g., ductal carcinoma in situ [DCIS], invasive ductal carcinoma [IDC], and invasive lobular carcinoma [ILC]).

In this study, we primarily used receptor status-based classification when such information was available. When studies provided only molecular status-based classification, we included HER2-enriched with HER2+ and basal-like with TNBC. If no subtype information was available, patients were labeled as Unknown. For more detailed information on each cohort, please refer to **Supplementary Table 1** and the original papers associated with the cohort identifiers.

Clinical information was extracted from each original publication, either from the released public data from the original publication, or from reaching out to the original authors. If both did not work out, we used the annotation from the TISCH2 database ^41^.

*EMTAB8107* ^42^: 14 treatment-naive BRCA patient samples were profiled using 5′ scRNA-seq with the 10X Genomics platform. A total of 44,024 high-quality cells were analyzed, comprising 16,235 tumor cells and 27,789 non-tumor cells. Among the 14 patients, 8 were TNBC, 4 were HER2+, 1 was luminal A-like, and 1 was luminal B-like. In terms of tumor-node-metastasis (TNM) staging, stages ranged from I-II. 13 samples were IDC while one was metaplastic carcinoma. The patient age range was 40–90 years. Cell type annotations were derived from the original study, and the gene expression count matrix and metadata were downloaded from https://lambrechtslab.sites.vib.be/en/pan-cancer-blueprint-tumour-microenvironment-0.

*GSE176078* ^43^: This cohort comprised 26 primary BRCA tumors, including 11 ER+, 5 HER2+, and 10 TNBC cases. Samples were profiled using scRNA-seq with the Chromium platform (10X Genomics). The gene expression matrix of 100,064 cells, including 24,489 tumor cells and 75,575 non-tumor cells, was downloaded from the Gene Expression Omnibus (GEO) under accession GSE176078. Among the 26 patient samples, 5 had previously received chemotherapy or targeted therapy, while 21 were treatment-naïve. Among the five patients receiving prior treatment, one received adjuvant chemotherapy (AC), one received neoadjuvant FEC-D, one received neoadjuvant AC four times and paclitaxel three times, and one received neoadjuvant AC four times and paclitaxel one time. The patient age range was 35–88 years. Grades ranged from II-III, and 22 samples were IDC, two were ILC, and two were MBC. Cell type annotations were derived from the original study.

*Bassez_CohortA* and *Bassez_CohortB* ^44^: Both cohorts were part of the same study ^44^, a single-center, open-label, non-randomized phase 0 trial (BioKey) (NCT03197389). The study included patients with non-metastatic, operable, newly diagnosed primary invasive breast carcinoma, histologically confirmed as ER−/PR− or ER+, with a primary tumor diameter >1 cm. In both cohorts, a single dose of 200 mg pembrolizumab (commercial name Keytruda, which is an anti-PD1 antibody) was administered before surgery in a window-of-opportunity setting. Fresh tumor tissue was collected from patients before pembrolizumab administration (via needle biopsy) and 7–15 days (9 ± 2) after administration (via surgical resection).

Bassez_CohortA involved 31 treatment-naive patients, including 13 TNBC, 3 HER2+, and 15 ER+ BRCA cases. A total of 175,942 high-quality cells were analyzed, comprising 55,432 tumor cells and 120,510 non-tumor cells. Among the 31 patients, 5, 23, and 3 were classified as TNM stage I, II, and III, respectively. The patient age range was 30–90 years. 27 samples were IBC-NST, one sample was ILC-ST, one was carcinoma with apocrine differentiation, and one was metaplastic carcinoma. 12 patients were pre-menopausal, and 19 patients were post-menopausal. Bassez_CohortB consisted of 11 patients with residual tumor on imaging after three months of neoadjuvant chemotherapy, including 6 TNBC, 1 HER2+, and 4 ER+ BRCA cases.

Chemotherapy was combined with anti-HER2 therapy for HER2+ tumors. A total of 50,693 high-quality cells were analyzed, comprising 10,453 tumor cells and 40,240 non-tumor cells. Among the 11 patients, 3, 7, and 1 were classified as TNM stage I, II, and IV, respectively. The patient age range was 30–80 years.

For both cohorts, single-cell transcriptomics were measured using 5′ scRNA-seq with the 10X Genomics platform. The gene expression count matrix and metadata were downloaded from https://lambrechtslab.sites.vib.be/en/single-cell.

*Wu* ^45^: This cohort comprised primary breast tumors collected from 5 TNBC patients. Among these, 4 patients were treatment-naive, and 1 patient had previously received adjuvant chemotherapy. All samples were grade III and one TNBC was metaplastic. Stage information was unspecified. Fresh tissue samples were processed using the Chromium platform (10× Genomics) for single-cell capture, followed by sequencing on the NextSeq 500 (Illumina) with the Chromium Single Cell 3’ v2 Library, Gel Bead and Multiplex Kit, and Chip Kit (10× Genomics). A total of 24,271 high-quality cells were analyzed, including 4,709 tumor cells and 19,562 non-tumor cells. Across these cells, 28,118 genes were detected. The patient age range was 35–73 years. The gene expression count matrix and cell type annotations were obtained from the original authors.

*GSE161529* ^46^: This large cohort included normal tissue, preneoplastic heterozygous BRCA1 mutation (BRCA1+/–) tissue, and BRCA cancer tissues from various subtypes. Among the 55 study subjects, 18 were non-cancer, 4 had preneoplastic BRCA1+/– tissue, 8 were TNBC, 6 were HER2+, 1 was PR+, and 18 were ER+ BRCA patients. Fresh tissue samples were subjected to 3′ scRNA-seq using the 10X Genomics Chromium platform. A total of 332,168 high-quality cells were analyzed, comprising 126,940 tumor cells and 205,228 non-tumor cells. The age ranges for the non-cancer, preneoplastic BRCA1+/–, and BRCA cancer patients were 19–49, 30–43, and 25–84 years, respectively. Among the 33 BRCA patients, 2, 7, and 24 were classified as TNM stage I, II, and III, respectively. All patients spanned grades I-III, and 2 patients were male. The gene expression count matrix and cell type annotations were derived from the TISCH2 database^41^.

*SRP114962* ^47^: This cohort comprised 8 pre- and post-NAC TNBC samples. SnRNA-seq was performed on fresh-frozen TNBC tissue samples, sequenced at 100 paired-end or 76 paired-end cycles on the HiSeq4000 system (Illumina). A total of 2,472 high-quality cells were analyzed, including 2,375 tumor cells and 97 non-tumor cells. Patients were with an age range of 39-58 years. Four patients were treatment responders, and four patients were treatment non-responders. Seven samples were IDC while one was unknown. The gene expression count matrix and cell type annotations were derived from the TISCH2 database ^41^.

*GSE140819* ^48^: This cohort included six metastatic BRCA patients with metastases to sites such as lymph nodes and liver. There were two lymph node metastases, three liver metastases, and two brain metastases. Four fresh tissue samples were analyzed using 3′ scRNA-seq with the 10X Genomics platform, while three frozen tissue samples were analyzed using 3′ snRNA-seq. For one patient, paired snRNA-seq and scRNA-seq data were obtained from two samples. Cancer subtype information for these patients was unavailable. A total of 63,908 high-quality cells were analyzed, comprising 45,121 tumor cells and 18,787 non-tumor cells. The gene expression matrix and cell annotation metadata were downloaded from the GEO under accession number GSE140819. Patient age information was not publicly available for this cohort.

*GSE148673* ^49^: This cohort included 6 treatment-naive BRCA patients, comprising 5 TNBC cases and 1 DCIS case. Fresh tumor tissue samples were obtained from MD Anderson Cancer Center. 3′ scRNA-seq was performed using the 10X Genomics platform. A total of 10,359 high-quality cells were analyzed, including 5,454 tumor cells and 4,905 non-tumor cells. The gene expression count matrix and cell type annotations were derived from the TISCH2 database ^41^. Patient age and tumor stage information was not publicly available for this cohort.

*GSE150660* ^50^: This cohort included 3 brain metastasis patients, comprising 2 with IDC and 1 with ILC. ScRNA-seq was performed using the 10X Genomics Chromium system with the Single Cell 3’ Library and Gel Bead Kit V2 (catalog no. 120234). A total of 10,605 high-quality cells were analyzed, including 1,843 tumor cells and 8,762 non-tumor cells. The patient age range was 26–88 years. The gene expression count matrix and cell type annotations were derived from the TISCH2 database ^41^. We did not use the expression count matrix from GEO because it was not accompanied by cell type annotations.

*GSE143423* ^51^: This cohort included 1 brain metastasis TNBC patient. Freshly resected tumor tissue was obtained from brain lesions of a treatment-naive TNBC patient. 3′ scRNA-seq was performed using the 10X Genomics platform. A total of 4,375 high-quality cells were analyzed, comprising 4,099 tumor cells and 276 non-tumor cells. The TNBC patient was 44 years old. The gene expression count matrix and cell type annotations were derived from the TISCH2 database^41^.

*GSE138536* ^52^: This cohort included 8 BRCA patients, comprising 6 ER+ and 2 TNBC cases. Four patients were classified as grade II, and four as grade III. Six samples were IDC and two samples were ILC. Samples were collected from the primary tumor site during surgical resection at Stanford Hospital and City of Hope National Medical Center. Cells were sequenced on the NextSeq 500 (Illumina) platform using the Smart-seq2 protocol, generating 2×76 or 2×151 bp paired-end reads. A total of 1,902 high-quality cells were analyzed, including 438 tumor cells and 1,464 non-tumor cells. The gene expression matrix and cell annotation metadata were downloaded from the GEO under accession number GSE138536. Patient age information was not publicly available for this cohort.

*GSE118389* ^53^: This cohort included 6 primary TNBC patients, with 1 patient classified as grade II and 5 patients as grade III. TNM stage was unspecified. Single-cell RNA sequencing was performed on 1,534 cells isolated from six fresh TNBC tumors. Among these, 1,112 of 1,189 cells were reliably classified into non-epithelial (n = 244) and epithelial (n = 868) cell types. Single sorted cells were sequenced using the Smart-seq2 platform in 38-bp paired-end mode on the NextSeq 500. The patient age range was 42–64 years. The gene expression matrix and cell annotation metadata were downloaded from the GEO under accession number GSE118389.

*GSE75688* ^54^: This cohort included 11 primary BRCA patients, comprising 2 ER+, 4 HER2+, and 5 TNBC cases. Ten were treatment-naïve, while one was treated with NAC and Herceptin. Two patients also had lymph node metastases (BC03 and BC07) sequenced. All surgical samples, except one, were obtained from chemotherapy-naive patients. The samples were processed using Fluidigm C1-based single-cell RNA sequencing combined with the SMARTer Ultra Low RNA Kit for full-length transcriptome amplification. Sequencing was performed in 100-bp paired-end mode on the HiSeq2500 platform. A total of 515 high-quality cells were analyzed, including 317 tumor cells and 198 non-tumor cells. The gene expression matrix and cell annotation metadata were downloaded from the GEO under accession number GSE75688. Patient age and tumor stage information was not publicly available for this cohort.

*GSE158724* ^55^: This cohort included 34 primary ER+ BRCA patients, all post-menopausal. SnRNA-seq was performed on serial biopsies collected from these patients, who were treated with neoadjuvant endocrine therapy (letrozole) with or without a CDK4/6 inhibitor (ribociclib). Samples were all taken at baseline (no treatment), and then either Arm A: letrozole + placebo, letrozole, Arm B: ribociclib 600mg (intermittent), or Arm C: letrozole + ribociclib 400mg (continuous). Within Arm A, N=11 patients, 6 responders and 5 non-responders. Within Arm B, N=12 patients, 6 responders and 6 non-responders. Within Arm C, N=11 pateints, 4 responders and 7 non-responders. Biopsies were obtained at three time points: beginning of treatment, 14 days after starting treatment, and after 180 days. Nuclei were isolated from the samples, and 3’ snRNA-seq was conducted using either the 10X Genomics or iCELL8 platforms. For the 10X Genomics-based sequencing, a total of 110,568 high-quality tumor cells were obtained. For the iCELL8-based sequencing, a total of 804 high-quality tumor cells were obtained. The gene expression matrix and cell annotation metadata were downloaded from the GEO under accession number GSE158724. While non-tumor cells were mentioned in the original study, the GEO repository did not provide data for non-tumor cells. Patient age and tumor stage information was not publicly available for this cohort.

#### In-house cohorts

*In-house 1*: This cohort included 82 primary, treatment-naïve tumors from women (ages 25-91) with BRCA, aged 25 to 91, across a range of molecular subtypes (41 luminal A, 9 luminal B, 6 HER2+, and 26 triple-negative). 8 participants were diagnosed at clinical stage I, 50 at stage II, 18 at stage III, and 1 at stage IV. Study participants were recruited from University of Maryland [NCI-Maryland BRCA studies (1993-2003 and 2012-2016), Aga Khan University Hospital, Nairobi, Kenya (AKUHN), and the AIC Kijabe Hospital, Kijabe, Kenya (both 2019-2021)]. All participants provided written informed consent prior to study enrollment and provided biospecimens at the time of surgery. Research pertaining to the NCI-Maryland studies was approved by the University of Maryland Institutional Review Board for the participating institutions (University of Maryland Medical Center and four surrounding hospitals in the Baltimore, Maryland, area) and by the National Institutes of Health Office for Human Research Protections, as previously described ^56,57^. The AKUHN and AIC Kijabe studies were approved by the Research and Ethics Committees at Aga Khan University Hospital, Nairobi (Ref: 2018/REC-80) and AIC Kijabe Hospital (KH IERC-02718/2019). A research license was obtained from the National Commission for Science and Technology (NACOSTI/P/24/33420) and a material transfer permit obtained from the Ministry of Health (MOH/ADM/1/1/81) in Kenya and followed recognized ethical guidelines as defined by the Declaration of Helsinki and the U.S. Common Rule. Samples were processed as frozen specimens, from which nuclei were isolated for 3’ snRNA-seq using the 10X Genomics platform. A total of 292,458 high-quality cells were analyzed, comprising 163,419 tumor cells and 129,039 non-tumor cells. Malignant and non-malignant cells were distinguished based on DNA copy number aberrations (aneuploidy) through CopyKAT ^49^. Among the patients, 66 samples have more than 500 tumor cells. This dataset, created by co-authors A.R.H., H.L., Sh.S., F.M., G.L.G., and S.A., has not yet been published and made publicly available and is thus described as “in-house”.

*In-house 2*: This cohort was part of a neoadjuvant clinical trial (NCT03366844) investigating the combination of pembro and focal radiotherapy (RT) in patients with TNBC. Among the 34 TNBC patients with a complete set of evaluable biopsies, 23 (67.6%) responded to treatment, achieving either a pathological complete response (pCR) with a Residual Cancer Burden (RCB) score of 0 (17/23) or showing minimal residual disease with an RCB score of 1 (6/23), as assessed by a breast pathologist at surgery. Ultrasound-guided biopsies were collected prior to the first cycle of pembro, three weeks after the first cycle of pembro, and three weeks after the second cycle of pembro and focal RT. Samples from these serial biopsies, obtained from 35 primary TNBC patients, were subjected to 5’ scRNA-seq using the 10X Genomics platform. A total of 514,495 high-quality cells were analyzed, comprising 50,185 tumor cells and 464,310 non-tumor cells. The patient age range was 33–94 years, with 1, 31, and 3 patients in clinical stages I, II, and III, respectively. The gene expression matrix and cell annotation metadata have been deposited in the GEO under accession number GSE246613 ^22^. We describe this cohort as “in-house” because S.V.R.K. made it available before it was deposited in GEO.

### Protein and RNA expression analysis of human normal tissues and cell types

#### RNA expression

scRNA-seq data from 31 human tissues were obtained from the Human Protein Atlas (HPA) database ^58^ (available at: https://www.proteinatlas.org/humanproteome/single+cell/single+cell+type/data#datasets). The dataset includes read counts per gene and cell, along with detailed annotations for tissue type and cell type. The 31 tissues analyzed encompass adipose tissue, bone marrow, brain, breast, bronchus, colon, endometrium, esophagus, eye, fallopian tube, heart muscle, kidney, liver, lung, lymph node, ovary, pancreas, peripheral blood mononuclear cells (PBMCs), placenta, prostate, rectum, salivary gland, skeletal muscle, skin, small intestine, spleen, stomach, testis, thymus, tongue, and vascular tissue.

Metadata, including data sources, cell counts, total read counts, and PubMed IDs, were compiled and tabularized in the provided link. The dataset comprises a total of 689,601 normal cells, providing a comprehensive resource for evaluating gene expression across diverse normal human tissues.

#### Protein expression

Protein expression data for 45 human tissues were obtained from the HPA. The expression profiles were generated using IHC on tissue microarrays. The dataset includes tissue names, annotated cell types, and expression levels categorized as “No,” “Low,” “Medium,” or “High,” in tabular form at the provided link. The data correspond to HPA version 24.0 and Ensembl version 109. The protein expression data is not at single-cell resolution.

#### Grouping and harmonization of tissues and cell types

To address discrepancies in tissue type and cell type annotations between RNA and protein expression datasets and to reduce the complexity of cell type classifications (e.g., over 100 distinct cell types), we performed a systematic classification of tissues and cell types.

Grouped Tissues: The harmonized tissue categories included heart, brain, hepatobiliary, lung, urinary, pancreas, bone marrow, integumentary, gastrointestinal, female reproductive, male reproductive, lymphoid, oral, visual, skeletal/smooth muscle, conducting airways, connective, endocrine (only for protein expression data), and vascular and PBMCs (only for RNA expression data).

Grouped Cell Types: The harmonized cell type categories included heart muscle, neuronal, alveolar, liver epithelial, urinary epithelial, hematopoietic, integumentary, endothelial/vascular, gastrointestinal epithelial, female reproductive, male reproductive, immune, visual, skeletal/smooth muscle, connective/stromal, endocrine, general/glandular epithelial, glial, unspecified, and undifferentiated and mixed (only for RNA expression data).

A detailed mapping between the grouped tissues/cell types and the original tissues/cell types is provided in **Supplementary Table 5**.

#### Equivalent expression of logic-gated gene circuits

To derive an equivalent expression for multi-gate circuits, comparable to single-gene expression, we employed the following translation methods. For an AND gate, the equivalent expression was defined as the minimum expression level of the two genes. For an OR gate, the equivalent expression was determined as the maximum expression level of the two genes. For a NOT gate, the equivalent expression was calculated as the negated expression of the gene.

In the context of protein expression, the original expression levels were categorized into “No,” “Low,” “Medium,” and “High” in the HPA data, which were numerically coded as 0, 1, 2, and 3, respectively. The NOT-gated equivalent expression was derived by subtracting the numerically coded expression of the protein from 3, the highest expression value. However, since the protein expression data is not at single-cell resolution, the equivalent expression of multi-gene circuits may represent an overestimation. For instance, if both gene A and gene B are expressed as “High” in a tissue, the expression of ’A & B’ is also classified as “High” under our rule. This approximation may not reflect the true expression at the single-cell level, as gene A and gene B could be highly expressed in different cells, resulting in an actual ’A & B’ expression of “Low” or even “No” in any single cell.

For scRNA-seq expression, we utilized log1p normalized counts (a common approach in scRNA-seq analysis, where raw gene expression counts for each cell were first normalized to a total of 10,000 counts per cell and then transformed using the natural logarithm of the normalized counts plus one), with expression values of 0, 1, 2, 3 (and >3) labeled as “No,” “Low,” “Medium,” and “High,” respectively. The NOT-gated equivalent expression was calculated in the same manner as for protein expression, ensuring consistency across both data types.

### Harmonization of data from different cohorts

To generate a unified UMAP visualization comprising all tumor cells from the 17 cohorts, we integrated the sc/snRNA-seq datasets using Harmony ^59^. This integration ensured that batch effects were minimized, allowing for a cohesive analysis across diverse datasets.

To prepare the input data for *LogiCAR designer*, the expression matrices—for both tumor cells from cancer patients and normal cells from 31 tissues of healthy donors—were filtered to include only cell surfaceome genes. The original expression values (raw or normalized counts) were binarized using a stringent threshold of 0, where any expression value greater than 0 was considered “expressed” and numerically set to 1. This approach resulted in uniform binarized sc/snRNA-seq matrices across all cohorts, intentionally disregarding potential batch effects that could introduce systematic differences in numeric expression values.

### Enumerating designs and circuits

In this study, we considered all three logic gates—AND, OR, and NOT—simultaneously for any N = 2, 3, 4, or 5 gene inputs. This led to an exponential growth in the number of unique designs. For example, even when treating two genes A and B as interchangeable (i.e., considering A & B and B & A as the same design), there were still six unique 2-gene designs: A & B, A | B, A & !B, A | !B, !A & !B, and !A | !B. For 3-gene, 4-gene, and 5-gene designs, the number of unique designs increased to 20, 70, and 252, respectively. In general, the number of designs that are not equivalent to another design by symmetry for N-gene input variables is given by the binomial coefficient C(2N, N), which represents the total number of ways to choose N items from 2N distinct items.

To reduce computational complexity and save coding and running time, we implemented a strategy to minimize the number of unique designs. We doubled the input surfaceome gene list by treating A and !A as two distinct genes, effectively eliminating the NOT gates from the designs. This simplification reduced the number of unique 2-gene designs to two: ‘A & B’ and ‘A | B’. For 3 genes, there were four unique designs: ‘(A & B) & C’, ‘(A | B) & C’, ‘(A & B) | C’, and ‘(A | B) | C’.

For designs involving four or more genes, we employed a recursive strategy from computer science. For instance, a 4-gene design could be constructed by combining a 3-gene design and a 1-gene design or by combining two 2-gene designs. Given that there are 1, 2, and 4 unique designs for 1, 2 and 3 genes, respectively, and two logic gates (AND or OR) to combine them, there are at most 4 × 2 × 1 + 2 × 2 × 2 = 16 potential designs for 4 genes. However, some of these designs may be duplicates by symmetry.

To eliminate duplicates, we utilized the concept of a truth table from computer science. In the truth table, each row represents a design, and each column represents a unique valuation possibility for the genes in the design. For example, for genes A, B, C, and D, the possible valuations range from (A, B, C, D) = (0, 0, 0, 0) to (1, 1, 1, 1). Each entry in the truth table matrix (i, j) corresponds to the calculated result (0 or 1) of the circuit in row i for the gene valuations in column j. By comparing rows in the truth table, we identified and removed duplicate designs, where two rows had identical values across all columns. This process yielded 12 unique designs for 4-gene circuits and 40 designs for 5-gene circuits.

All unique designs were hard coded into the *LogiCAR designer* algorithm and are provided in Supplementary Table 6.

### The LogiCAR designer algorithm

#### Input data

The *LogiCAR designer* algorithm requires two primary inputs:

● A binarized single-cell gene expression matrix of tumor cells, where 0 indicates “not expressed” and 1 indicates “expressed.”
● A binarized single-cell gene expression matrix of healthy normal cells, using the same binary encoding.

These matrices serve as the foundation for evaluating candidate CAR circuits.

#### The scoring function

The algorithm aims to maximize tumor cell-targeting efficacy while maintaining a normal cell-sparing safety threshold. Specifically:

● Tumor-targeting efficacy (%) is calculated as the percentage of tumor cells expressing the candidate CAR (either a single-gene or a multi-gene circuit) among all tumor cells.
● Normal cell-sparing safety (%) is calculated as the percentage of normal cells that do not express the candidate CAR among all normal cells.

For each possible circuit c, both efficacy and safety are evaluated. The final objective score is computed as follows:

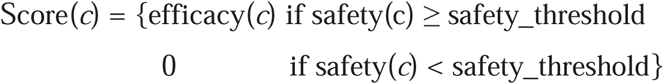

#### The iteration of optimal solutions

*LogiCAR designer* employs a genetic algorithm, which mimics the natural evolution process *in silico.* The algorithm operates as follows:

1. Initialization: Randomly generate a population of N-gene circuits with a population size of M. This initial population represents potential solutions and is denoted as generation 0 (G0).
2. Evaluation: Calculate the objective scores for all M circuits in the current generation.
3. Selection: Select two circuits from the current generation, with the probability of selection proportional to their objective scores. Higher-scoring circuits have a greater chance of being selected, but lower-scoring circuits still retain a small probability, mimicking natural selection.
4. Crossover: Perform crossover on the selected circuits by randomly setting a breakpoint and exchanging genes after the breakpoint to create two new circuits.
5. Mutation: Randomly mutate genes in the new circuits by substituting them with random genes from the surfaceome gene pool. Mutation occurs with a probability pmut, simulating natural genetic mutations.
6. Population Update: Repeat steps 3–5 to generate a new population of size M.
7. Diversity Enhancement: Every Ggap generations, replace the lowest-scoring Rrep percentage of circuits with randomly generated circuits. This mimics natural population perturbations, such as invasions, to increase diversity.
8. Termination: Repeat steps 2–7 until a stop criterion is met.
9. Output: Return the L circuits with the highest objective scores.

#### Implementation

*LogiCAR designer* is implemented in Python using the DEAP (Distributed Evolutionary Algorithms in Python) framework (version 1.4.1) and Python 3.9. The script leverages NumPy (version 1.26.4) for optimized Boolean operations during expression pattern evaluation. For each candidate circuit, the pipeline evaluates expression patterns, computes confusion matrix elements, calculates efficacy and safety scores, and stores results for post-processing.

To mitigate the risk of the genetic algorithm converging to local optima, 10 independent runs with different random seeds were performed for each design. The best solutions from all runs were merged and further selected.

Regarding the stopping criteria, for a given N-gene design, the program terminates if either of the following conditions is met:

⍰ No better solutions (higher objective scores) emerge in the most recent 200 generations.
⍰ The total number of generations reaches G_max_.

where G_max_ is 100, 500, 800, and 1000 for N =2, 3, 4, and 5, respectively. The other parameters were set as follows:

⍰ Population size (M): 1000
⍰ Mutation probability (p_mut_): 0.2
⍰ Generational replacement interval (G_gap_): 10
⍰ Replacement percentage (R_rep_): 10%

### Computational time complexity

For a specific N-gene design, the number of possible ways to assign the gene rages from binomial coefficient C(S, N), where S is the total number of candidate genes, if all genes are interchangeable by symmetry, to the product S × (S-1) × … (S - N +1) if no gene pairs are interchangeable by symmetry. As noted earlier, there are C(2N, N) unique designs. Therefore, the time complexity for an exhaustive search (Tes) of all N-gene circuits is:

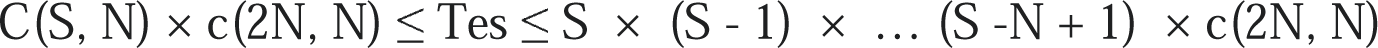

For *LogiCAR designer*, the time complexity (Tld) is determined by the number of unique logic-gated designs C(N), the number of circuits tested in each generation M, the total number of generations G, and the number of random seeds used for repeated runs R:

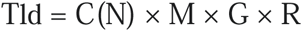

where the values of M and R were fixed at 1000 and 10, respectively. For N = 2, 3, 4, 5, the values of C(N) were 2, 4, 12, 40, as described earlier, and the values of G were capped at 200, 400, 800, and 1000, respectively.

### The curation of cell membrane surface genes

Cell membrane surface genes were curated from The In Silico Human Surfaceome database ^60^ (http://wlab.ethz.ch/surfaceome/). A false positive rate threshold of 5% (SURFY scores > 0.5818) was applied. This list was further augmented with targets tested in solid tumor clinical CAR trials that were not already included (see below).

The final curated list comprised 2,758 unique cell surface genes (**Supplementary Table 7**). Among these, 2,725 were measured in the union set of all the BRCA clinical cohorts, and 2,724 were measured in HPA normal cells. A total of 2,716 surfaceome genes were shared between the tumor cell matrix and the HPA normal cell expression matrix and were used as input for LogiCAR Designer. Consequently, all newly identified candidate CARs were derived from this final set of 2,716 genes (**Supplementary Table 7**).

Throughout the manuscript, targets are referred to by their official Human Genome Nomenclature Committee (HGNC) gene symbols in italic font (e.g., *TACSTD2*) rather than the names of the encoded proteins in non-italic font (e.g., TROP2).

### The curation of solid-tumor CAR targets from existing or planned clinical trials

Solid-tumor clinical CAR targets were obtained using Trialtrove ^61^ on January 23, 2024 (**Supplementary Table 2**). Hence, some newer trials such as NCT06347068 were omitted. Access to Trialtrove requires a license. For each clinical trial, the target was identified based on the description of the drug product, supplemented by manual curation to resolve gene aliases and, in a few cases, by queries submitted via the “Ask the Analyst” feature of Trialtrove.

### The curation of BRCA CAR circuits from previous computational methods

Previously identified solid tumor CAR circuits were obtained from two studies. The first study, by ^12^, predicted circuits from bulk RNA expression data. The identified circuits were downloaded from https://antigen.princeton.edu/ (**Supplementary Table 2**). For this study, the top 20 circuits for Breast Carcinoma, ranked by Combined Score, were extracted and used.

The second study, by ^16^, identified circuits based on single-cell expression data. The circuits were extracted from https://static-content.springer.com/esm/art%3A10.1038%2Fs41587-023-01686-y/MediaObjects/41587_2023_1686_MOESM7_ESM.xlsx (**Supplementary Table 2**). The top 20 circuits for BRCA, ranked by the mean Expression-Cell Fraction (ECF) of tumor cells, were selected using a safety criterion of <10% ECF in normal cells, consistent with the threshold set in this study. However, it was noted that many of these circuits included non-cell membrane surface genes, such as *ATP5MC1, KRT8, KRT18*, and *KRT19*. After consulting with the original authors, we confirmed this was an error in their original work. Consequently, circuits containing non-surface genes were excluded from our comparison.

### Tumor-targeting efficacy grouping

Patient-level tumor-targeting efficacy was classified using three thresholds designed to align with clinical criteria for minimal pathological response, partial response, and complete response. In clinical practice, a 10% reduction in the maximal tumor radius is considered a minimal pathological response (e.g., ^25^), while a 30% reduction is classified as a partial response according to the RECIST (Response Evaluation Criteria in Solid Tumors) guideline ^26^. The absence of detectable tumor cell signals is defined as a complete response.

To simplify the analysis, tumors were assumed to be solid spheres. Under this assumption, 10% or 30% reductions in radius correspond to 27% or 66% reduction in volume (or tumor cells), respectively. Based on this, we established 27% and 66% tumor-targeting efficacy as thresholds for candidate CAR circuits. To account for numerical inaccuracies, we further defined a 99% tumor-targeting efficacy threshold to approximate complete tumor clearance.

## Statistical analyses

All statistical analyses were conducted using Python (v3.9) or R (v4.1). No assumptions were made regarding the normality of data distributions, and all statistical tests were performed as two-tailed analyses.

## Data availability

All the publicly available cohorts are in **Supplementary Table 1** with links to the data. The de-identified gene expression matrix of in-house cohort 1 will be made publicly available upon acceptance of this and a companion work. The gene expression matrix of in-house cohort 2 has been made publicly available at GEO246613. The Harmony integrated data of all cohorts will be made publicly available online at Zenodo.

## Acknowledgements

This work is supported in part by the Intramural Research Program of the National Institutes of Health, NCI. This work utilized the computational resources of the NIH HPC Biowulf cluster (http://hpc.nih.gov).

## Author contributions

S.M., E.R., and A.A.S. conceived and designed the study. T-G.C. and S.M. developed the algorithm. S.M., T-G.C., S.R.D., B.W., and A.A.S., acquired, analyzed or interpreted the data. T-G.C., S.M., A.A.S., C-P.D., and E.R drafted the manuscript. A.R.H, H.L., Sh. S., F.M., G.L.G, S.A. collected in-house data set 1 and advised on analysis of it. A.M. and S. R. V. K. advised on the analysis of in-house data set 2, which was published during the study. P.S.R., A.S., and Sa.S. advised on some analyses. All authors critically revised the manuscript for important intellectual content. E.R. supervised the study.

