## Supplementary Figures for "Single-cell-guided identification of logic-gated antigen combinations for designing effective and safe CAR therapy"

**Supplementary information**

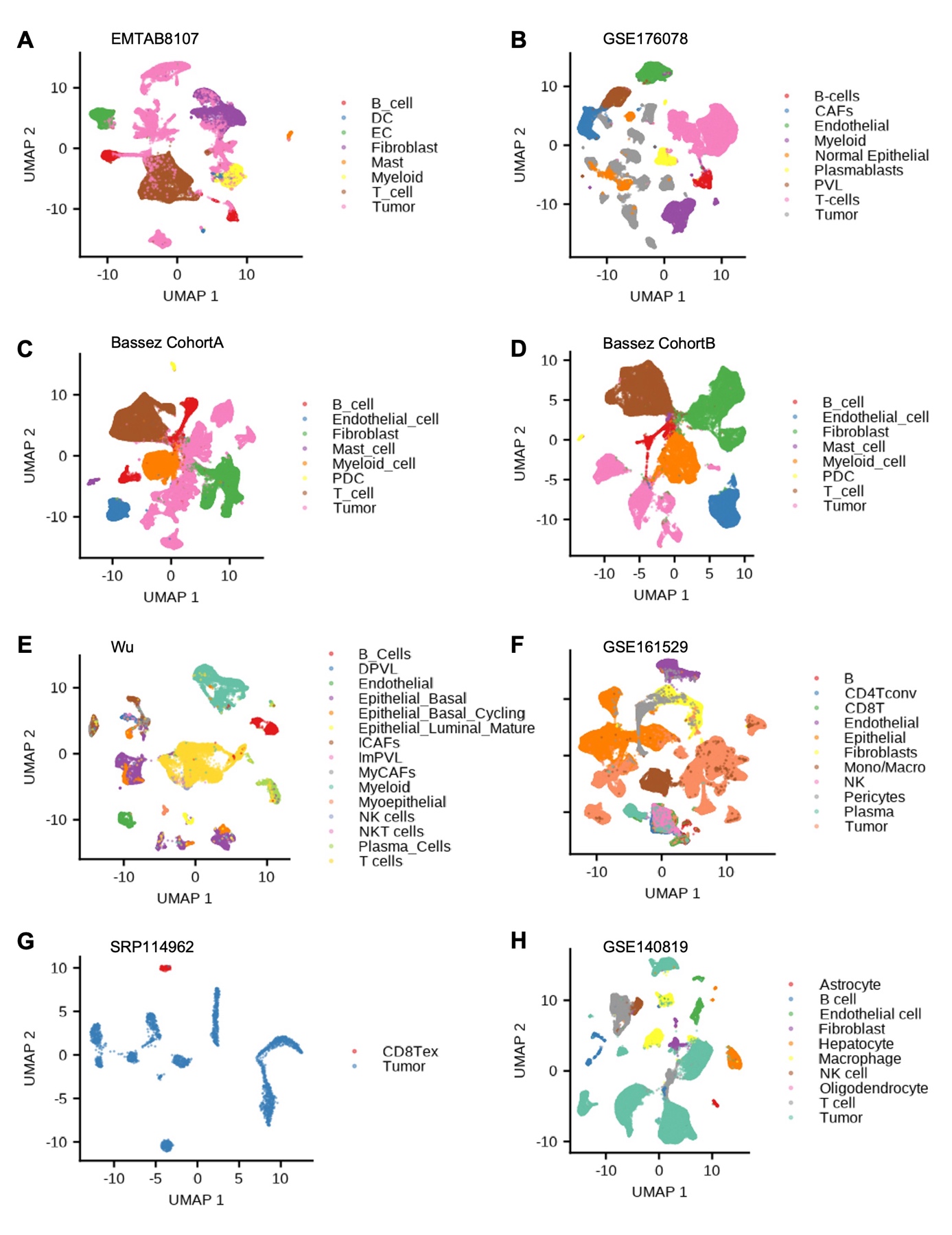

**Supplementary Figure 1. (continues on the next page)**

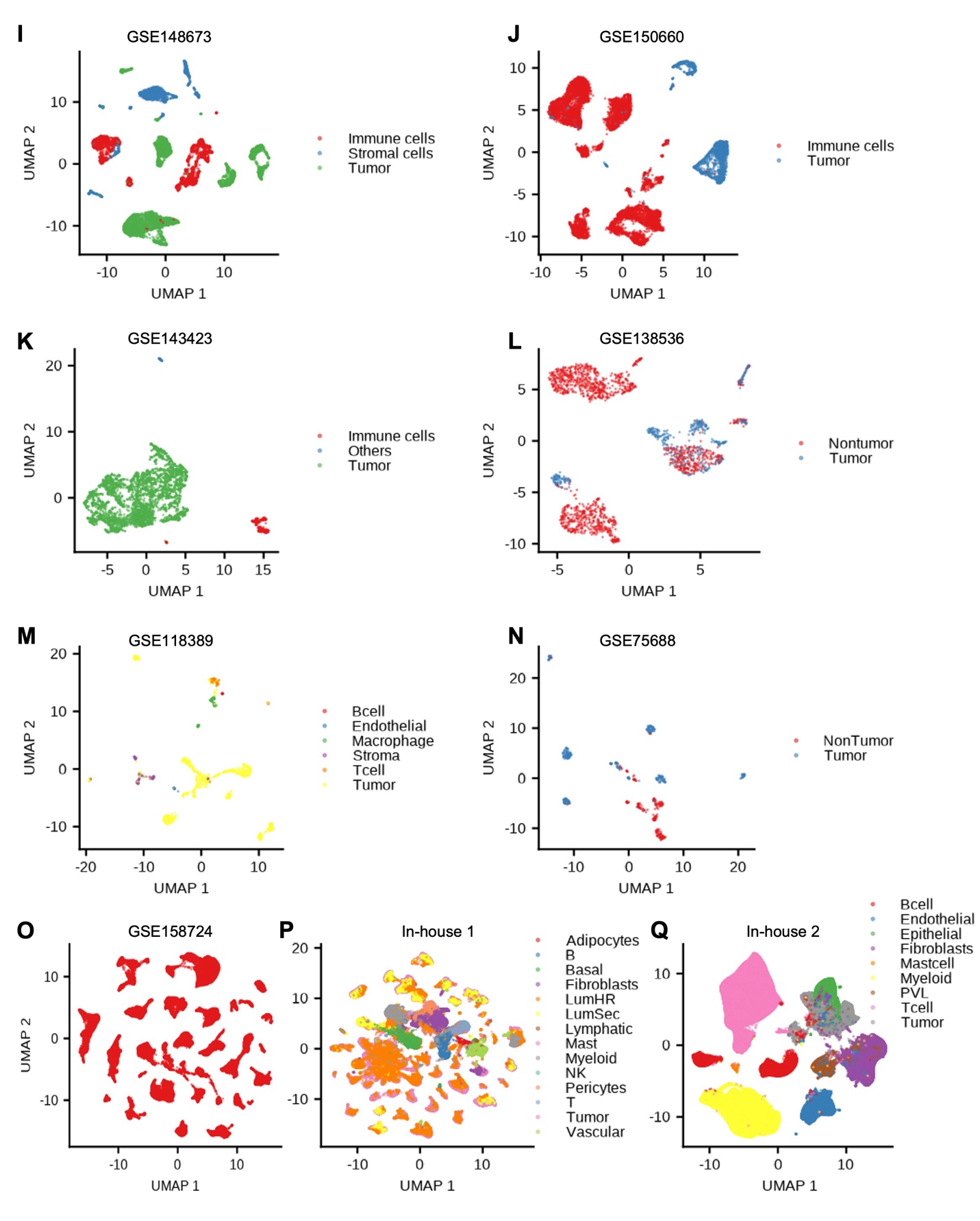

**Supplementary Figure 1. UMAP Representation of All Cell Types Measured in the Tumor Microenvironment of the 17 BRCA Clinical Cohorts. (related to Figure 1)**

Note that in cohort GSE158724, only tumor cells were measured.

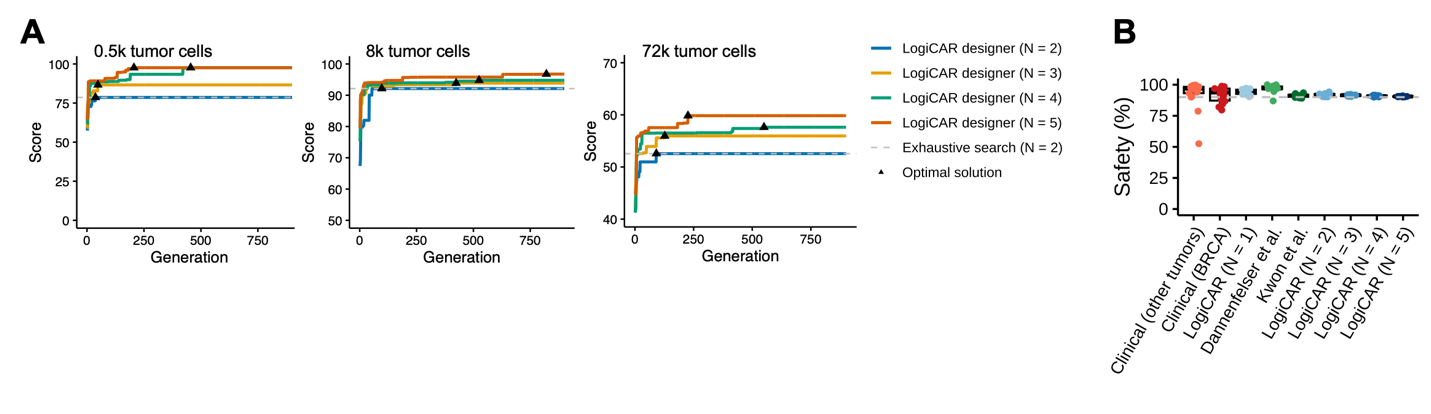

**Supplementary Figure 2. Design Safe and Effective Logic-Gated Combinatorial CAR Targets for Breast Cancer. (related to Figure 2)**

**A**. The number of generations (right panel of Fig. 1C) needed for convergence of *LogiCAR designer* to achieve the optimal solution with gene numbers in combinations of N = 2, 3, 4, 5, and with the input number of tumor cells set to 500 (left), ~8,000 (middle), and ~74,000 (right). The dashed lines represent ground truth from exhaustive search for N = 2. Triangles mark the earliest generations of iterations in *LogiCAR designer* that identified the optimal solution.

**B**. The safety score distribution of CAR target combinations in different groups in C. Safety was calculated from the randomly sampled 31k normal cells from the HPA curation.

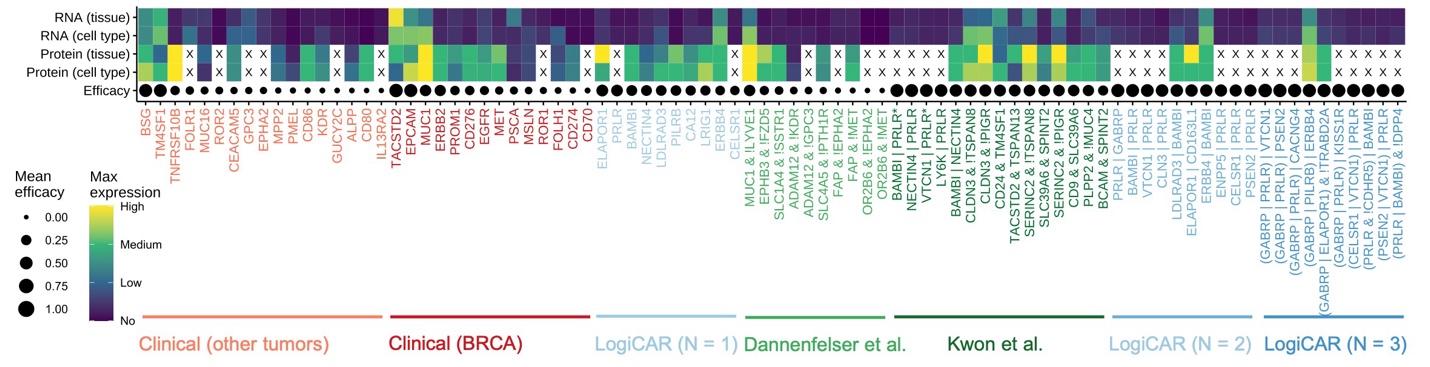

**Supplementary Figure 3. Characterizing Safety Landscape of Potential CAR Targets. (related to Figure 3)**

A summary of the RNA and protein expression levels of potential CAR target combinations across normal tissues and cell types, respectively. RNA or protein expression is represented as the maximal mean expression across all normal tissues or cell types in **Fig. 3A**. Efficacy is reported as the mean efficacy across all 17 cohorts.

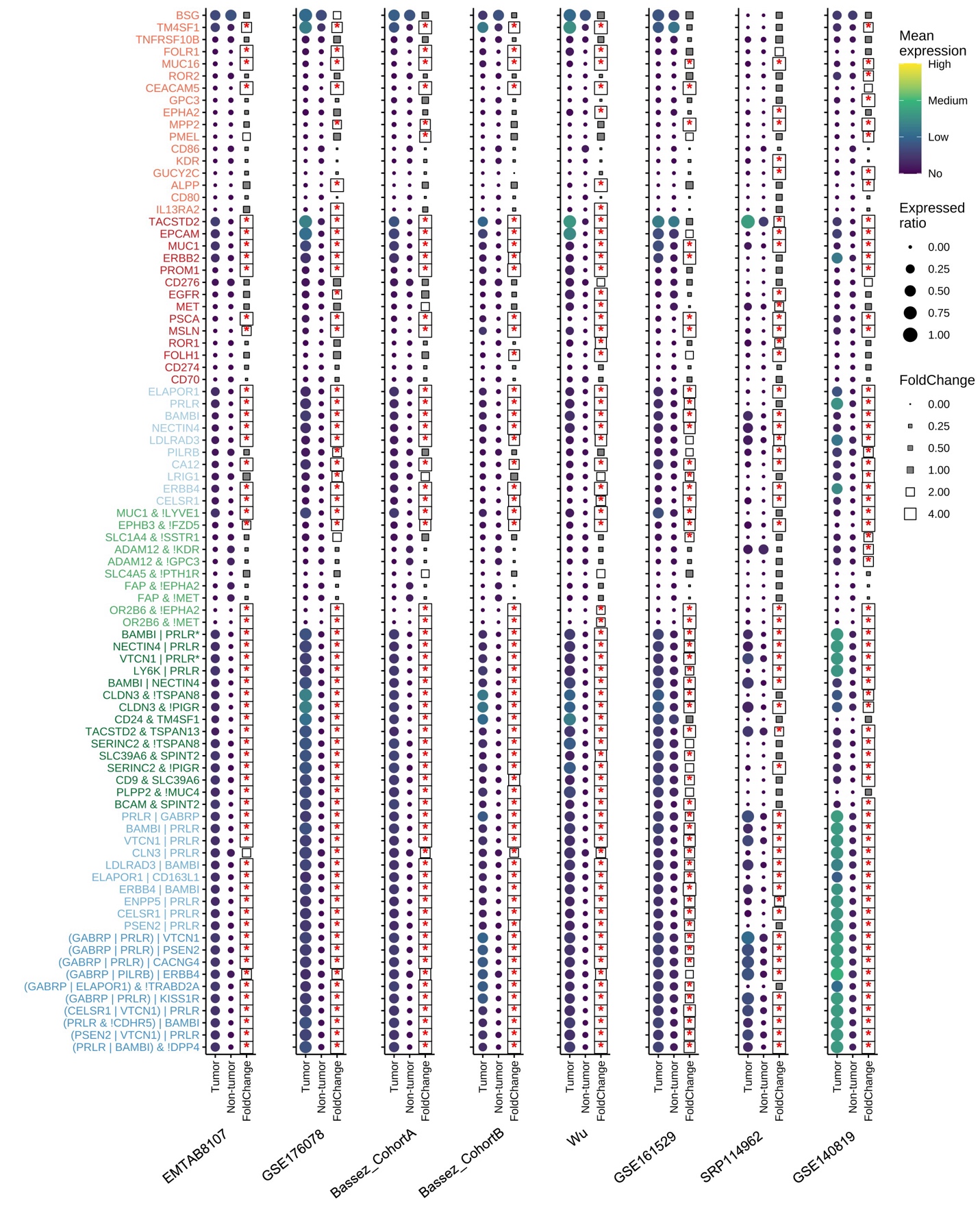

**Supplementary Figure 4. (continues on the next page)**

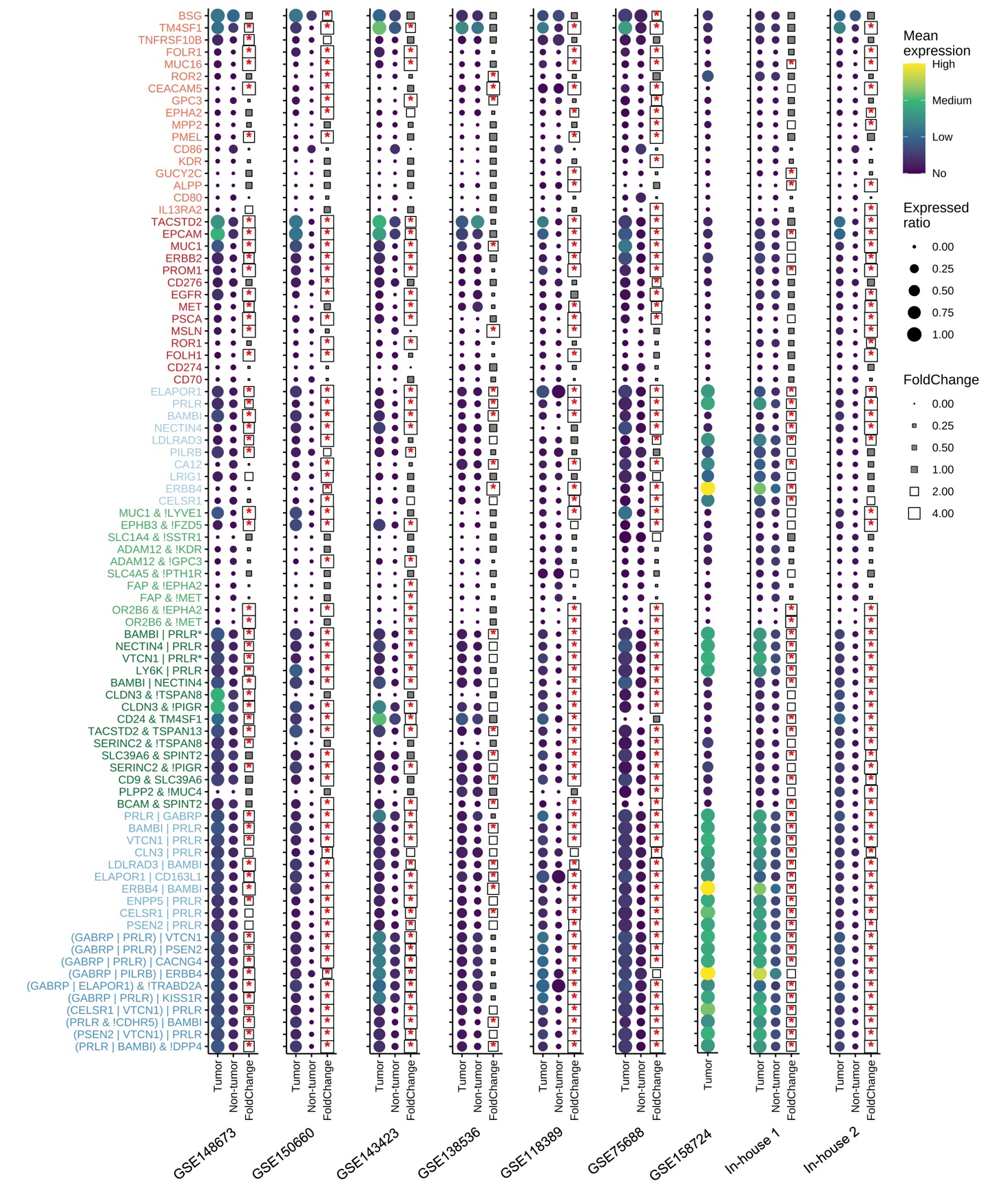

**Supplementary Figure 4. Comparing Expression Pattern of Potential CAR Targets among Tumor versus Non-tumor Cells in the Tumor Microenvironment. (related to Figure 3)**

RNA-level expression of LogiCAR-designed circuits and single-gene CARs in tumor cells and non-tumor cells in the tumor microenvironment (TME) across 17 BRCA datasets. The fold changes are log-normalized single-cell RNA expression between tumor cells and non-tumor cells for each CAR target or circuit. Statistical significance was determined using a two-tailed Wilcoxon test, with p-values adjusted for multiple comparisons using the Bonferroni method. Data points with an adjusted p-value < 0.05 and a fold change > 2 are marked with an asterisk (*). Note that in cohort GSE158724, only tumor cells were measured.

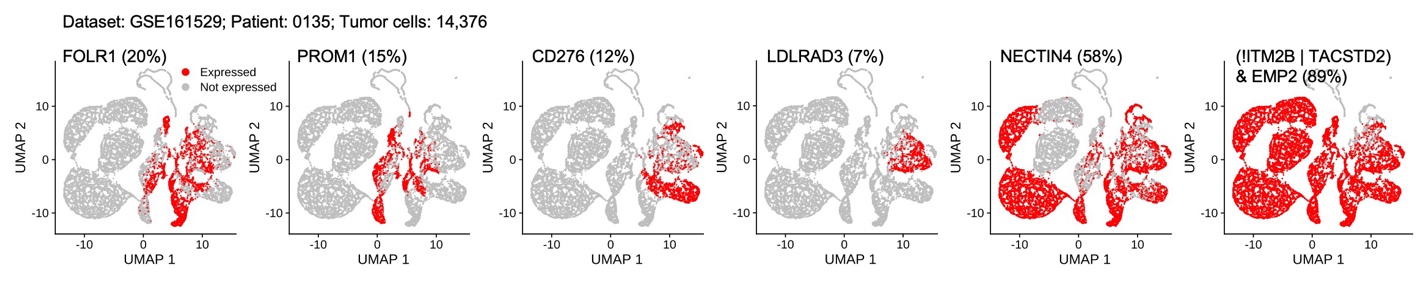

**Supplementary Figure 5. Personalized CAR Target Design for a TNBC Patient to Address Intratumoral Heterogeneity. (related to Figure 5)**

Personalized CAR targets effectively address the tumor heterogeneity challenge. The UMAP was created based on the input of a union of surface genes shown. “Expressed” indicates that a tumor cell has an expression level of the corresponding CAR higher than zero. The proportion of tumor cells expressing the corresponding CAR target or combination is shown alongside the CAR target or combination.
